# Museomics analyses inform about *Channichthys* icefish species diversity

**DOI:** 10.1101/2024.09.25.615019

**Authors:** Benedicte Garmann-Aarhus, Ekaterina Nikolaeva, Thomas Desvignes, Nicolas Straube, Michael Matschiner

## Abstract

The rapid diversification of notothenioid fishes in the waters surrounding the Antarctic continent is a prime example of the process of adaptive radiation. Within around 10 million years, Antarctic notothenioids have diversified into over 100 species with a broad range of lifestyles and ecological adaptations. However, the exact number of species within this radiation has long been unclear. Particularly challenging is the taxonomy of the genus *Channichthys*, for which between one and nine species have been recognized by different authors. The putative species from this genus are known from a limited number of representative specimens, of which most were sampled decades ago. Here, we investigated the mitochondrial genomes of museum specimens representing the four recently recognized species Unicorn Icefish (*C. rhinoceratus*), Red Icefish (*C. rugosus*), Sailfish Pike (*C. velifer*), and Charcoal Icefish (*C. panticapaei*), complemented by morphological analyses. All analyzed specimens were collected in the 1960s and 1970s and fixed in formaldehyde, and their DNA has thus been heavily degraded. Applying ancient-DNA protocols for DNA extraction and single-stranded library preparation, we were nevertheless able to obtain sufficient endogenous DNA to reconstruct the mitochondrial genomes of one specimen of each species. These mitochondrial genome sequences were nearly identical for the three specimens assigned to Unicorn Icefish, Red Icefish, and Sailfish Pike, while greater mitochondrial divergence was observed for the Charcoal Icefish specimens. We discuss possible explanations of the contrast between these molecular results and the recognizable morphological variation found among the four species, and recommend that at least the Charcoal Icefish be included the list of valid icefish and notothenioid species.

## 1 Introduction

Examples of the process of adaptive radiation – the rapid proliferation of species diversity as the result of ecological opportunity (Schluter, 2000; Simpson, 1953) – are known almost exclusively from terrestrial and freshwater environments (Grant & Grant, 2008; Lerner, Meyer, James, Hofreiter, & Fleischer, 2011; Losos, 2009; Ronco et al., 2021). One remarkable example of this process in the marine system, however, is found in the freezing waters of Antarctica. Here, fishes of the suborder Notothenioidei – the Antarctic members of which are collectively termed “Cryonotothenioidea” (Near et al., 2015) – have diversified into over 100 species (Eastman & Eakin, 2021), characterized by an incomparable array of phenotypes and lifestyles (Bista et al., 2023; Clarke & Johnston, 1996; Eastman, 2005; Matschiner et al., 2015; Near et al., 2012, 2018).

The diversity of cryonotothenioids is classified into five families, of which icefishes (family Channichthyidae) are arguably the most iconic. Not only are these fishes the only known vertebrates to lack hemoglobin (Desvignes, Bista, Herrera, Landes, & Postlethwait, 2023; Eastman, 1993; Ruud, 1954), they also carry numerous adaptations, such as antifreeze glycoproteins and enlarged hearts and gills (DeVries & Wohlschlag, 1969; Rankin & Tuurala, 1998; Sidell & O’Brien, 2006; Tuta, Acierno, & Agnisola, 1991), making them well suited for the highly oxygenated, freezing waters of the Southern Ocean. Along with their range of adaptations, the family of icefishes has evolved very recently, with genome-based age estimates placing their origin within the past 5 million years (Bista et al., 2023). Despite their young age, they today include at least 16 species classified into eleven genera (Eastman & Eakin, 2021).

The numbers of species within each of the eleven genera of the family Channichthyidae are in most cases uncontroversial. One genus, however, has been a topic of discussion for years – the genus *Channichthys*, distributed exclusively around the sub-Antarctic Kerguelen and Heard Islands (Duhamel, Gasco, & Davaine, 2005). First described in 1844 by Richardson, the genus has expanded from being monotypic to including as many as nine classified species (Ahyong et al., 2024; Froese & Pauly, 2024). These are the Unicorn Icefish (*C. rhinoceratus* Richardson, 1844), the Red Icefish (*C. rugosus* Regan, 1913), the Sailfish Pike (*C. velifer* Meisner, 1974), the Aelita Icefish (*Channichthys aelitae* Shandikov, 1995b), the Big-eyed Icefish (*C. bospori* Shandikov, 1995b), the Pygmy Icefish (*C. irinae* Shandikov, 1995b), the Charcoal Icefish (*C. panticapaei* Shandikov, 1995a), the Green Icefish (*C. mithridatis* Shandikov, 2008), and the Robust Icefish (*C. richardsoni* Shandikov, 2011). However, in recent years, these classifications have become subject to scrutiny due to new morphological analyses (Nikolaeva, 2019, 2020). These analyses led to the synonymization of several of the nine species, leaving only the Unicorn Icefish (*C. rhinoceratus*), the Red Icefish (*C. rugosus*), the Sailfish Pike (*C. velifer*), and the Charcoal Icefish (*C. panticapaei*), recognized by Eschmeyer’s Catalog of Fishes (Fricke, Eschmeyer, & van der Laan, 2024). These four species were found to be distinguishable by phenotypic traits such as their gill rakers, coloration, fin height, fin-ray count, and head and snout shape (Nikolaeva, 2019, 2020, 2021, 2024; Nikolaeva & Balushkin, 2019).

Genetic analyses corroborating this classification have long been lacking, due to the difficulty of obtaining fresh samples from the Kerguelen-Heard Plateau and the technical challenges of extracting suitable DNA from older, museum specimens. A particular obstacle to molecular analyses of museum specimens has been that such specimens were commonly fixed in formaldehyde, a substance that breaks and alters DNA strands (Straube, Lyra, et al., 2021). Thus, it was only with the development of ancient-DNA protocols facilitating the extraction and sequencing of DNA from museum specimens (Gansauge, Aximu-Petri, Nagel, & Meyer, 2020; Gansauge et al., 2017; Straube, Lyra, et al., 2021), that the first molecular insights were gained into the classification of *Channichthys* icefishes. In 2022, Muschick et al. were able to isolate and sequence sufficient DNA from a formaldehyde-fixed specimen of the Red Icefish (*C. rugosus*) that allowed the assembly of nearly its entire mitochondrial genome (NCBI accession PP319402.1). The comparison of this mitochondrial genome with one that was publicly available for the Unicorn Icefish (*C. rhinoceratus*; NCBI accession NC 057120) revealed great similarity between these two mitochondrial genomes, leading the authors to suggest that the two species might in fact be one and the same (Muschick et al., 2022).

Here, we expand on the investigation by Muschick et al. (2022) with additional museum specimens, representing all of the four recently recognized *Channichthys* species (Nikolaeva, 2020). We apply ancient-DNA protocols to tissue samples of these specimens to extract DNA for molecular analyses based on Illumina sequencing, and complement these analyses with a new morphological assessment of species differences. Of twelve specimens sampled around the Kerguelen Islands in the late 1960s and early 1970s, one of each recognized species yielded sufficient suitable DNA to reconstruct its mitochondrial genome sequence. Our molecular analyses showed that not only the Unicorn Icefish (*C. rhinoceratus*) and the Red Icefish (*C. rugosus*) specimens have nearly identical mitochondrial genomes, but that this is also the case for the Sailfish Pike (*C. velifer*) despite its rather clear phenotypic differentiation. In contrast, the mitochondrial genome of the Charcoal Icefish (*C. panticapaei*) specimen was more distinct, with a degree of separation comparable to that found in a second pair of icefish congeners. We speculate that these results may indicate a smaller number of *Channichthys* species than previously assumed, that interspecific hybridization might have led to exchange of mitochondrial genomes, or that the genus might be in the process of a new and rapid radiation on the Kerguelen-Heard Plateau.

## 2 Materials and Methods

### 2.1 Sampling

Twelve *Channichthys* icefish specimens were selected for molecular analysis. All of these are part of the collection of the Zoological Institute of the Russian Academy of Sciences in Saint Petersburg, Russia, and were collected during cruises by Soviet and – in one case (ZIN 53007) – French scientists in the late 1960s and early 1970s. All specimens were fixed in formaldehyde after their capture, and were eventually transferred to 70% ethanol. They were assigned to the four recently recognized species following the updated identification key of Nikolaeva (2020).

### 2.2 DNA extraction and sequencing

DNA extraction, library preparation, and quality checks (qPCR, concentration measurement, automated electrophoresis) were performed at the molecular laboratory of the University Museum of Bergen. As far as possible, all laboratory procedures followed recommendations detailed in Fulton and Shapiro (2019), such as UV-C irradiation, strict decontamination protocols, and positive pressure HEPA filtration.

For all twelve specimens, DNA was extracted from muscle tissue (between 28 and 54 mg dry weight per sample), applying the same treatment to all samples, as well as two extraction negatives. DNA extraction followed the guanidine treatment described in Straube, Lyra, et al. (Appendix S1; 2021) which is based on and modified from Basler et al. (2017); Dabney et al. (2013); Rohland, Glocke, Aximu-Petri, and Meyer (2018); Rohland and Hofreiter (2007a, 2007b); Rohland, Siedel, and Hofreiter (2010); Sambrook and Russell (2001). Before amplification and simultaneous dual indexing of the single-stranded libraries, qPCR was used to determine the ideal individual number of amplification cycles. Final concentrations were measured in a Qubit dsDNA High-Sensitivity assay (Thermo Fisher Scientific), and fragment length distributions and maximum peak sizes were assessed with Bioanalyzer High Sensitivity DNA Analysis (Agilent). All libraries were used in shallow shotgun sequencing for initial data generation to check for the presence of target DNA (see 2.3). Initial sequencing was performed on an Illumina MiniSeq system at the University Museum of Bergen. Deeper sequencing for specimens identified to contain sufficient target DNA (ZIN 56275, ZIN 56538, and ZIN 56638) was performed on the Illumina NextSeq system at the University of Potsdam (Adaptive Genomics Group). Additional sequencing efforts were guided by initial mapping experiments to a mitochondrial reference genome (see 2.3). The numbers of reads mapping to this genome were then used to estimate the additional amount of mitochondrial reads necessary to allow for reconstructing the mitochondrial genome (see Straube, Preick, Naylor, and Hofreiter 2021).

All sequencing used Illumina single-end high output kits following Paijmans et al. (2017). For those samples for which additional sequencing was performed, the resulting sets of reads were combined. Similarly, for the specimen that had previously been sequenced in Muschick et al. (2022) (ZIN 56294), new sequencing data generated herein were merged with those generated previously. Illumina’s bcl2fastq2 tool (https://support.illumina.com/sequencing/sequencingsoftware/bcl2fastq-conversion-software.html) was used for demultiplexing.

### 2.3 Molecular analyses

Sequence reads were processed with fastp v.0.23.2 (Chen, Zhou, Chen, & Gu, 2018) to remove adapter sequences, the first base of the tail of forward reads, bases with Phred-scaled quality below 25 at both ends of reads, and bases in poly-G regions with a length greater than 10 bp. We then excluded reads if they carried more than 10 undetermined bases, if they were shorter than 20 bp, or if more than 40% of bases had a Phred-scaled quality below 20.

To verify the presence of endogenous DNA and estimate its quantity, we assigned each sequence read to a taxonomic group by mapping to the NCBI non-redundant (NR) sequence database (downloaded on 21 November 2023). This assignment used the BLASTX search algorithm as implemented in the program Diamond v.2.1.6 (Altschul, Gish, Miller, Myers, & Lipman, 1990; Buchfink, Xie, & Huson, 2015). The frequencies of different taxon assignments were then plotted with MEGAN v.6.25.3 (Huson et al., 2016).

To identify endogenous mitochondrial reads, we prepared a chimeric reference sequence from the mitochondrial genome of *Channichthys rhinoceratus* (NCBI accession NC 057120) and the nuclear genome of the closely related species *Pseudochaenichthys georgianus* (EBI accession GCF 902827115.1) (Bista et al., 2023). We mapped reads to this reference using the Paleomix (v.1.3.8) BAM pipeline (Schubert et al., 2014) with BWA MEM (Li & Durbin, 2010) selected as aligner software. The pipeline included the application of the program AdapterRemoval (Schubert, Lindgreen, & Orlando, 2016), excluding reads with a minimum length 25 bp or a mismatch rate greater than 3. Also as part of the Paleomix pipeline, we invoked the program mapDamage (Jónsson, Ginolhac, Schubert, Johnson, & Orlando, 2013) to assess DNA degradation. The resulting files in BAM format were used for variant calling with GATK’s Germline Short Variant Discovery (v.4.5.0.0) pipeline (McKenna et al., 2010), a method suitable for the identification of variants from haploid data such as mitochondrial sequences. We focused on the four specimens that yielded most mitochondrial reads per recognized species (ZIN 56638, ZIN 56294, ZIN 56275, and ZIN 56538) in this and the subsequent analyses. Following variant calling, the resulting variant dataset was filtered using bcftools v.1.18 (Li, 2011), considering only variants with Phred-scaled quality score above 30 and a read depth greater than 3. We further verified the identified mitochondrial variants through manual inspection with the Integrative Genome Viewer v.2.18.2 (Robinson et al., 2011), confirming that the single nucleotide polymorphisms (SNPs) identified were not artifacts from post mortem DNA damage (e.g., variants near the ends of individual reads). Based on the confirmed variant calls and the mitochondrial reference sequence, we reconstructed the full mitochondrial genome sequences for the four focal specimens, ensuring that sites that had insufficient coverage for variant calling were coded as missing data. We then annotated the mitochondrial genome sequences with the MitoAnnotator pipeline (Zhu, Sato, Sado, Miya, & Iwasaki, 2023).

For phylogenetic analyses, we generated an alignment with MAFFT v.7.520 (Katoh & Standley, 2013) from the four reconstructed mitochondrial genome sequences, together with the available reference sequence for *C. rhinoceratus* (NC 057120). Based on this alignment, we calculated the pairwise genetic distances among all five sequences, and performed maximum-likelihood phylogenetic inference with IQ-TREE v.2.2.2.7 (Minh et al., 2020). This inference included IQ-TREE’s substitution model selection and the application of 1,000 ultrafast bootstrap replicates (Minh, Nguyen, & von Haeseler, 2013).

To embed the *Channichthys* sequences within the broader context of notothenioid diversification, we downloaded mitochondrial genome sequences for 34 additional notothenioid species (Supplementary Table 1; e.g. Papetti et al. 2021). We extracted the sequences of the 13 mitochondrial protein-coding genes from these genomes as well as from the five *Channichthys* genomes. For each of these genes, we then aligned all 39 sequences, with MAFFT. From these alignments, we extracted all first, second, and third codon positions, and concatenated these into three codon-specific partitions. The mitochondrial phylogeny of the included notothenioid species was then reconstructed in a partitioned maximum-likelihood analysis with IQ-TREE that otherwise had the same settings as before.

Finally, we aimed to compare sequences of individual regions from the four reconstructed *Channichthys* mitochondrial genomes to publicly available sequences of orthologous regions, isolated from other specimens of the genus. The only mitochondrial marker with sufficient available sequences, however, was cytochrome c oxidase I (*mt-co1*), for which twelve *C. rhinoceratus* sequences were available (NCBI accessions JN640603–JN640614; Smith et al. 2012). We downloaded these twelve sequences and aligned these to those of the four reconstructed genomes and the reference, again using MAFFT. From this alignment, a haplotype genealogy graph was produced with the program Hapsolutely v.0.2.2 (Vences et al., 2024), selecting maximum parsimony as the method for tree inference and Fitchi (Matschiner, 2016) for the graph reconstruction.

### 2.4 Morphological analyses

We complemented the molecular analyses of the four *Channichthys* specimens ZIN 56638 (*C. rhinoceratus*), ZIN 56294 (*C. rugosus*), ZIN 56275 (*C. velifer*), and ZIN 56538 (*C. panticapaei*) with a detailed assessment of morphological variation within the genus. This assessment focused on the four above-named specimens, but also incorporated a broad set of further specimens from the Zoological Institute of the Russian Academy of Sciences, Saint-Petersburg (ZIN), the National Museum of Natural History, National Academy of Sciences of Ukraine, Kyiv (IZANU), and the Natural History Museum, London (BMNH). A total of 280 specimens were studied, including the holotypes of *Channichthys rugosus* and *C. panticapaei*.

- *Channichthys rhinoceratus*: 160 specimens with *SL* from 150 to 395 mm, sampled around Kerguelen Island at known depths from 140 to 420 m. These specimens included ZIN 53006, ZIN 55586, ZIN 55587, and ZIN 56631– 56644. For more details, see Nikolaeva (2020).
- *Channichthys rugosus*: 10 specimens with *SL* from 215 to 370 mm, sampled around Kerguelen Island at known depths from 64 to 120 m. These specimens included ZIN 53007 and ZIN 56291–56994, as well as the holotype BMNH 1876.3.23.4 with *SL* 370 mm. For more details, see Nikolaeva (2021).
- *Channichthys velifer*: 45 specimens with *SL* from 134 to 490 mm, sampled around Kerguelen Island at known depths from 130 to 157 m. These specimens included ZIN 53005, ZIN 54807, ZIN 56271, and ZIN 56273–56290. For more details, see Nikolaeva and Balushkin (2019).
- *Channichthys panticapaei*: 61 specimens with *SL* from 138 to 398 mm, sampled around Kerguelen Island at known depths from 80 to 380 m. These specimens included ZIN 56520–56538 as well as the holotype IZANU 5109 with *SL* 348 mm. For more details, see Nikolaeva (2019).

The morphological analyses used a protocol developed and optimized for the study of *Channichthys* icefishes (Balushkin & Spodareva, 2015; Nikolaeva, 2016, 2017, 2019, 2020, 2021; Nikolaeva & Balushkin, 2019). For each specimen, 50 different morphological characters and indices were recorded (10 morphometric characters, 19 meristic characters, and 21 indices; see section 3.3). The features of the gill apparatus, seismosensory system, skin granulation (the degree of its development), and coloration were also examined. Additionally, the axial skeleton was studied using a digital microfocus X-ray diagnostic unit “PRDU-02” (“ELTECH-MED”, Saint-Petersburg, Russia) in the Centre of Collective Use “TAXON” at the Zoological Institute of the Russian Academy of Sciences.

All measurements were made visually, using a caliper, or in the case of X-ray images, with the program PRDU, developed by the platform provider. All calculations were performed with Excel (Microsoft) and Statistica (TIBCO Software). Based on our morphological analysis, a revised diagnosis of the genus and the four recently recognized species was developed.

## 3 Results

### 3.1 DNA extraction and sequencing

The application of ancient-DNA protocols to the formaldehyde-fixed museum specimens of the genus *Channichthys* led to mixed results across the twelve specimens. The extracts had DNA concentrations between 1.43 ng/µl (ZIN 56273) and values below the Qubit High Sensitivity kit detection threshold of 0.05 ng/µl (ZIN 56538) (Table 2). Due to the sample-specific amplification step during library preparation, these libraries were more homogeneous in their post-amplification DNA concentrations, ranging from 3.4 to 19.7 ng/µl. Maximum peak sizes in the fragment length distributions were detected between 153 and 175 bp.

**Table 1.** *Channichthys* icefish specimens analyzed.

| Specimen | Species | Sampling date | Sampling location | Depth |
| --- | --- | --- | --- | --- |
| ZIN 56635 | *C. rhinoceratus* | 16 Mar 1969 | NA | 280 m |
| ZIN 56636 | *C. rhinoceratus* | 13 Jan 1970 | NA | NA |
| ZIN 56638 | *C. rhinoceratus* | 24 Dec 1970 | 48°04' S, 70°15' E | 145–150 m |
| ZIN 53007 | *C. rugosus* | Jan 1964 | NA | NA |
| ZIN 56294 | *C. rugosus* | 28 Jun 1974 | 48°34' S, 70°37' E | 115–120 m |
| ZIN 56292 | *C. rugosus* | 1967 | NA | NA |
| ZIN 56273 | *C. velifer* | 26 Jan 1970 | NA | NA |
| ZIN 56274 | *C. velifer* | 20 Feb 1970 | NA | NA |
| ZIN 56275 | *C. velifer* | 20 Feb 1970 | 48°58' S, 67°27' E | 195–207 m |
| ZIN 56536 | *C. panticapaei* | 24 Feb 1969 | NA | NA |
| ZIN 56537 | *C. panticapaei* | 16 Mar 1969 | NA | 280 m |
| ZIN 56538 | *C. panticapaei* | 26 Dec 1970 | 48°00' S, 70°32' E | 148 m |

Specimen ZIN 56294 was previously used for molecular analyses in Muschick et al. (2022). NA, not available.

**Table 2.** Summary of DNA extraction and Illumina MiniSeq sequencing results.

| Specimen | Species | Concentration (ng/μl) |  | MiniSeq read numbers |  |
| --- | --- | --- | --- | --- | --- |
|  |  | Extract | Library | Total | Mapping |
| ZIN 56635 | *C. rhinocerus* | 0.20 | 19.7 | 1,150,880 | 0 |
| ZIN 56636 | *C. rhinocerus* | 0.42 | 12.5 | 1,496,964 | 0 |
| ZIN 56638 | *C. rhinocerus* | 0.80 | 4.6 | 1,701,466 | 131 |
| ZIN 53007 | *C. rugosus* | 0.34 | 4.3 | 1,701,825 | 3 |
| ZIN 56294 | *C. rugosus* | 1.10 | 3.6 | 1,753,908 | 109 |
| ZIN 56292 | *C. rugosus* | 0.94 | 3.4 | 1,530,944 | 51 |
| ZIN 56273 | *C. velifer* | 1.43 | 6.9 | 2,051,860 | 16 |
| ZIN 56274 | *C. velifer* | 0.81 | 9.0 | 1,627,555 | 29 |
| ZIN 56275 | *C. velifer* | 0.84 | 6.5 | 1,164,653 | 1,079 |
| ZIN 56536 | *C. pantacapai* | 0.22 | 4.7 | 1,208,445 | 0 |
| ZIN 56537 | *C. pantacapai* | 0.36 | 11.8 | 915,238 | 1 |
| ZIN 56538 | *C. pantacapai* | NA<sup>1</sup> | 5.6 | 2,212,835 | 646 |

Concentrations were measured in a Qubit dsDNA High-Sensitivity assay. The numbers of reads mapping refers to those that map to the mitochondrial genome (using NC\_057120 as reference).
<sup>1</sup> Concentration too low for measurement with Qubit High-Sensitivity assay (notably, despite a comparatively large number of reads mapping to the mitochondrial genome).

Sequencing on the Illumina MiniSeq platform produced between 915,238 and 2,215,285 reads per specimen (Table 2). Initial mapping to the reference showed that only few of the libraries contained at least moderate numbers (*>* 100) of mitochondrial reads. Based on these results, the libraries for ZIN 56275, ZIN 56538, and ZIN 56638 were re-sequenced for additional data on the Illumina NextSeq platform. For the specimen that had previously been sequenced in Muschick et al. (2022) (ZIN 56294), new sequencing data generated herein were combined with those generated previously. Similarly, unpublished NexSeq sequencing data previously generated for specimen ZIN 56292 were also combined with the new MiniSeq data.

Assigning sequencing reads to taxonomic groups with Diamond and MEGAN revealed that most DNA libraries consisted primarily of non-endogenous DNA, dominated by bacterial (such as *Burkholderia*) sequences (Supplementary Figures 1-12). Human contamination seemed to be minimal, as only three specimens (ZIN 53007, ZIN 56292, and ZIN 56275) had few reads assigned to Catarrhini (Old World monkeys). Out of the twelve specimens, four contained reads assigned to Notothenioidei, and for each of these four specimens, those reads were present in substantial proportions: ZIN 56638 (*C. rhinoceratus*), ZIN 56294 (*C. rugosus*), ZIN 56275 (*C. velifer*), and ZIN 56538 (*C. panticapaei*) (Fig. 2). Thus, all further molecular analyses focused on these four specimens.

**Figure 1.**
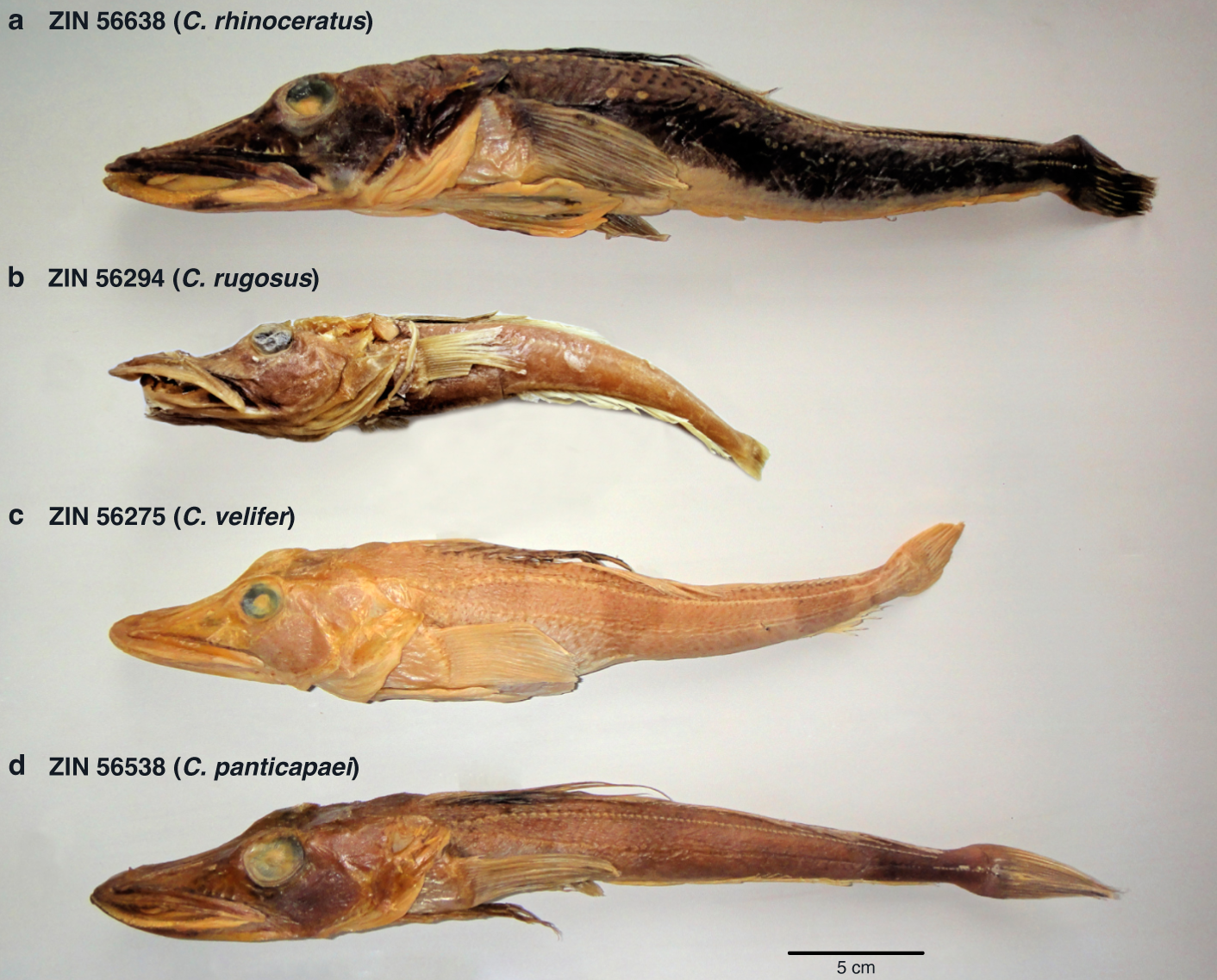
Specimens analyzed of the four recently recognized *Channichthys* species. (**a**) ZIN 56638 (*C. rhinoceratus*), (**b**) ZIN 56294 (*C. rugosus*), (**c**) ZIN 56275 (*C. velifer*), (**d**) ZIN 56538 (*C. panticapaei*).

**Figure 2.**
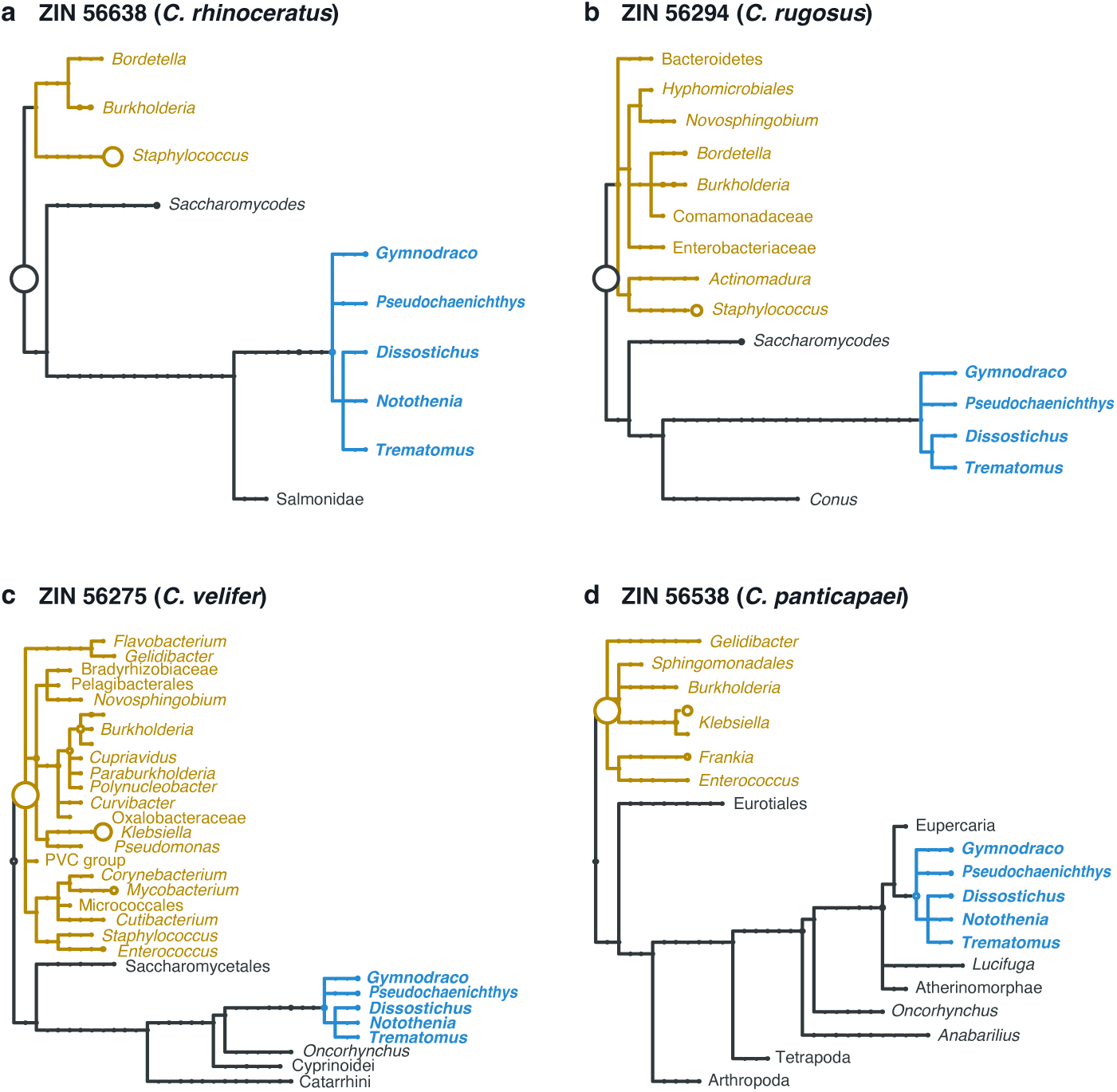
Taxonomic assignments of reads extracted from the four focal *Channichthys* specimens. Circles on nodes indicate frequencies of assignments. Bacterial taxa are colored in brown; notothenioid taxa are shown in blue. Assignments were made with the programs Diamond and MEGAN.

### 3.2 Molecular analysis

An assessment of DNA degradation with mapDamage showed elevated rates of C to T substitutions near the ends of reads, indicating deamination of cytosine residues to uracil. However, this deamination appeared to affect only the terminal 1–2 bp of each read (Supplementary Figures 13-16). For the four focal specimens ZIN 56638 (*C. rhinoceratus*), ZIN 56294 (*C. rugosus*), ZIN 56275 (*C. velifer*), and ZIN 56538 (*C. panticapaei*), Paleomix retained 11,711,034, 39,994,409, 20,801,663, and 35,235,998 reads, respectively. These sets of reads had mean lengths of 47.9, 35.9, 41.8, and 44.9 bp. Of the reads per specimen, 1,856, 10,873, 10,279, and 7,081 reads mapped to the mitochondrial genome of the *C. rhinoceratus* reference (NC 05712), respectively (Table 3). These numbers corresponded to 0.0158%, 0.0272%, 0.0494%, and 0.0201% of the retained reads per specimen.

**Table 3.**
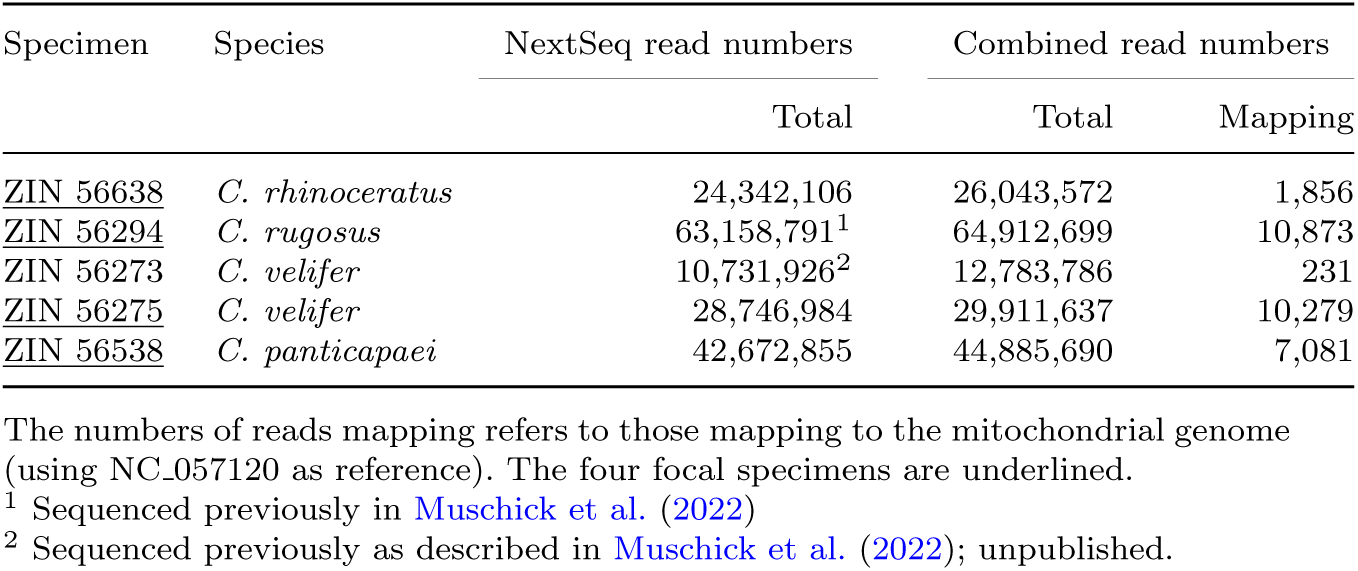
Summary of Illumina NextSeq sequencing results.

Analysis with GATK identified 140 variants, of which one was an indel – a deletion of one guanine residue at position 16,596 of the mitochondrial genome (within the D-loop / control region) in ZIN 56538 (*C. panticapaei*). The remaining 139 variants were SNPs. All variants had a Phred-scaled quality score greater than 30 and a coverage of at least 3 reads per specimen, and thus passed the filtering with bcftools. Manual inspection with Integrative Genome Viewer confirmed all 140 variants. Of the 139 SNPs, 119 were transitions and 20 were transversions, resulting in a transition/transversion ratio of 5.95. The number of C to T substitutions (22) was comparable to other transitions (A to G: 43; G to A: 24; T to C: 30). The mitochondrial genome sequence generated in this way for ZIN 56294 (*C. rugosus*) was nearly identical to that published by Muschick et al. (2022) for the same specimen (NCBI accession PP319402.1). The differences between the two versions were limited to the exact beginning and end positions of stretches of missing information (coded as “N”) in the D-loop region and for a single site in the same region (position 17,347) that was coded as “C” in the previous version of this genome, but now appeared to be more likely a “T”. The other three new mitochondrial genome sequences were uploaded to NCBI with accessions 3928 (ZIN 56275; *C. velifer*), 3929 (ZIN 56538; *C. panticapaei*), and 3930 (ZIN 56638; *C. rhinoceratus*). Annotation with the MitoAnnotator pipeline identified all 13 coding genes, the two rRNA genes, and the 22 tRNA genes in each of the reconstructed mitochondrial genomes.

Genetic distances were low for all pairs of specimens except those involving ZIN 56538 (*C. panticapaei*), with 13–34 changes among the mitochondrial genomes of ZIN 56638 (*C. rhinoceratus*), ZIN 56294 (*C. rugosus*), and ZIN 56275 (*C. velifer*) while 110–114 changes separated the mitochondrial genome of ZIN 56538 (*C. panticapaei*) from all other specimens and the reference (NC 057120; *C. rhinoceratus*) (Table 4). The phylogeny inferred with IQ-TREE from the five full mitochondrial genome sequences of *Channichthys* specimens (including the reference) illustrated this pattern with a long branch separating ZIN 56538 (*C. panticapaei*) from a cluster formed by the other four specimens (Fig. 3a).

**Figure 3.**
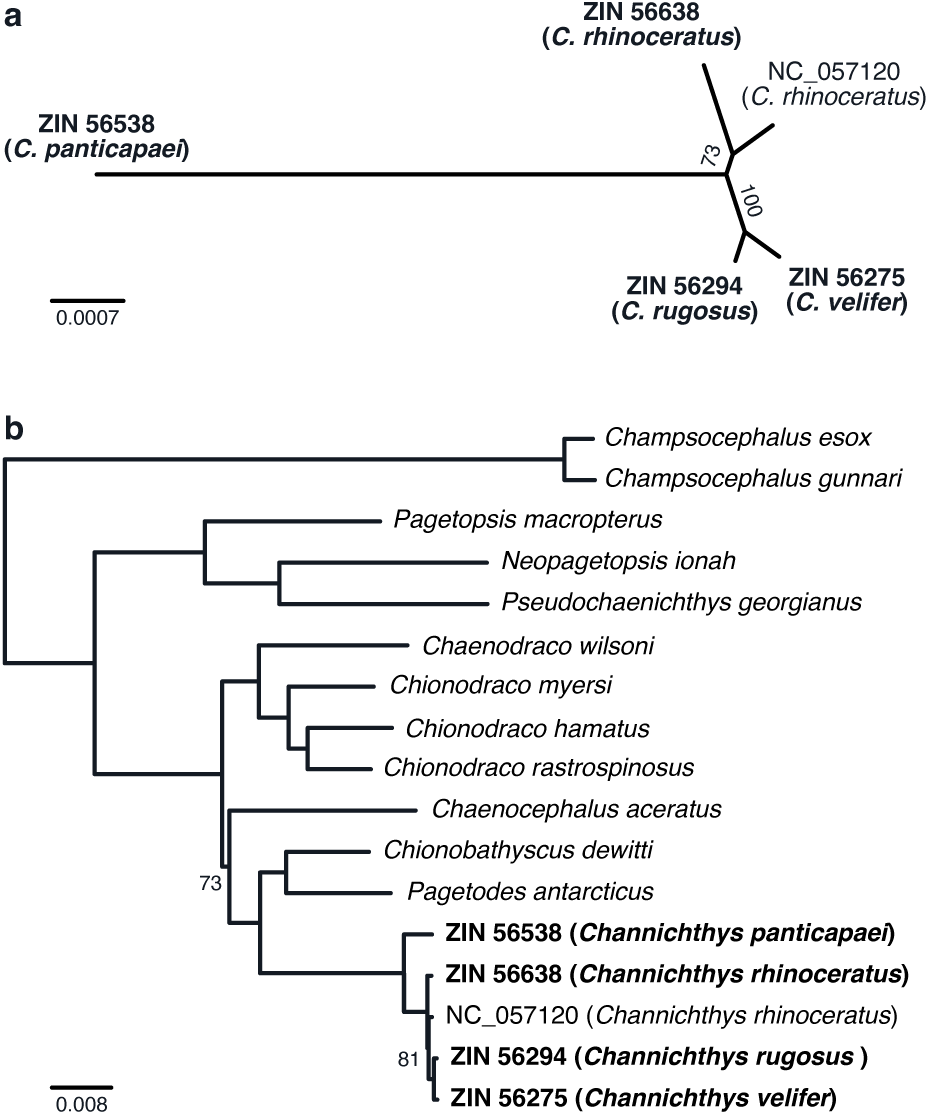
Maximum-likelihood phylogenies inferred for the genus *Channichthys* and the family *Channichthyidae*. **a** Unrooted phylogeny based on the four newly reconstructed mitochondrial *Channichthys* genomes (ZIN 56638, ZIN 56294, ZIN 56275, and ZIN 56538) and the *C. rhinoceratus* reference (NC 057120). **b** Rooted phylogeny for all species of Channichthyidae with available mitochondrial genomes, based on an alignment of coding sequences. In **a** and **b**, branch lengths mark substitutions per site, and labels on branches indicate bootstrap support. In **b**, bootstrap support is only shown for nodes that did not receive full support (BS *<* 100). Newly reconstructed mitochondrial genomes are marked in bold font.

**Table 4.**
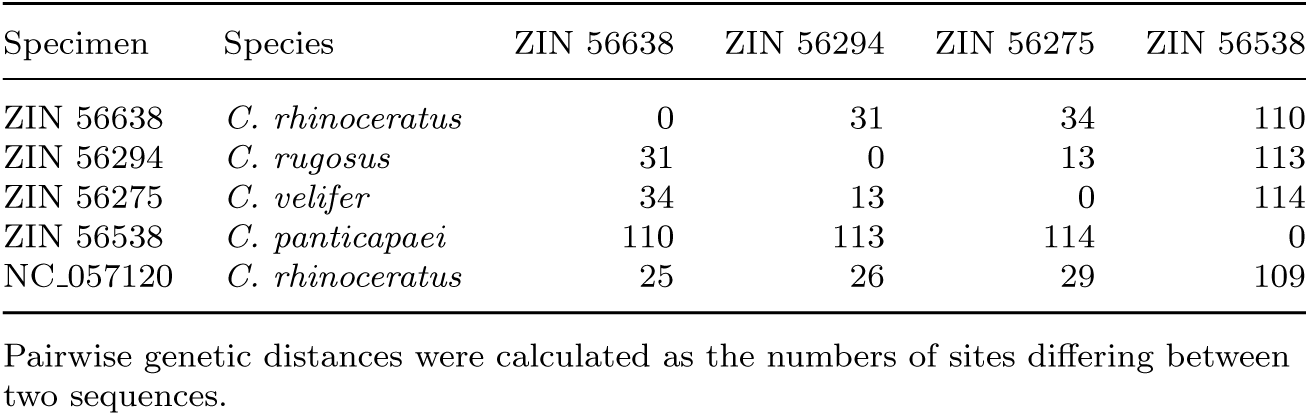
Pairwise genetic distances.

The partitioned alignment of mitochondrial coding sequences for the four newly reconstructed mitochondrial genomes, the *C. rhinoceratus* reference, and 34 further notothenioid species had a length of 11,460 bp with a high level of completeness (99.4%). Maximum-likelihood phylogenetic analysis based on this alignment produced a tree that was highly congruent with previous studies of notothenioid relationships (Colombo, Damerau, Hanel, Salzburger, & Matschiner, 2015; Dornburg, Federman, Lamb, Jones, & Near, 2017; Near et al., 2012, 2018). Focusing on Channichthyidae, we recovered the genus *Champsocephalus* (with the two representatives *C. esox* and *C. gunnari*) as the sister clade to all other included icefishes (Fig. 3b), with full bootstrap support (BS: 100). The remaining members of the family formed three strongly supported clades (BS: 100); one combining the genera *Pagetopsis*, *Neopagetopsis*, and *Pseudochaenichthys*, a second grouping *Chaenodraco* and *Chionodraco*, and a third clustering *Chionobathyscus*, *Pagetodes*, and *Channichthys*. All of these three clades have also been supported by nuclear data (Dornburg et al., 2017; Near et al., 2018). In line with phylogenies based on nuclear data, we also recovered the genus *Chaenocephalus* as the sister to the third of the three clades, albeit with weak support from our mitochondrial data (BS: 73). Within *Channichthys*, this phylogeny based on mitochondrial coding sequences (Fig. 3b) slightly differed from the one based on full mitochondrial genomes (Fig. 3a), as the two sequences assigned to *C. rhinoceratus* (ZIN 56538 and NC 057120) formed a clade in the latter, but not the former. However, the relationships among these two sequences were not strongly supported in either of the two phylogenies (BS: 73, 81). Concordant with the calculated pairwise genetic distances, the branches connecting the four *Channichthys* sequences assigned to *C. rhinoceratus*, *C. rugosus*, and *C. velifer* were shorter (below 0.0008 substitutions per site) than any other branches on the tree. The branch leading to ZIN 56538 (*C. panticapaei*) was still short compared to most other terminal branches, but comparable to those connecting the two congeners within the genus *Champsocephalus* (*C. esox* and *C. gunnari*; branch lengths ranging from 0.0038 to 0.0041 substitutions per site). In comparison, the three congeners within the genus *Chionodraco* had branch lengths that were more than twice as long (at least 0.0085 substitutions per site).

Cytochrome c oxidase I (*mt-co1*) haplotypes varied little among the newly reconstructed *Channichthys* genomes, the *C. rhinoceratus* mitochondrial reference genome (NC 057120), and twelve additional *mt-co1* sequences available from NCBI (which had all been assigned to *C. rhinoceratus*) (Fig. 4). A central haplotype was shared by 13 specimens, including those of the three specimens ZIN 56275 (*C. velifer*), ZIN 56294 (*C. rugosus*), and ZIN 56638 (*C. rhinoceratus*). This haplotype differed by one single mutation from that found in the *C. rhinoceratus* reference (NC 057120) and from two further haplotypes found in other sequences assigned to *C. rhinoceratus*. In contrast, the haplotype of specimen ZIN 56538 (*C. panticapaei*) was the most divergent, separated by five substitutions from the central haplotype.

**Figure 4.**
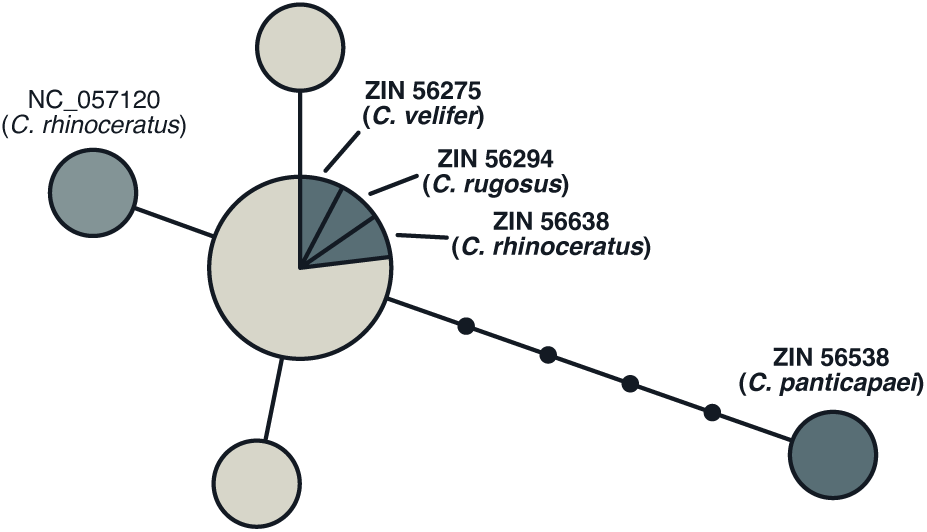
Haplotype genealogy graph for *mt-co1* sequences of the genus *Channichthys*. The three specimens ZIN 56275 (*C. velifer*), ZIN 56294 (*C. rugosus*), and ZIN 56638 (*C. rhinoceratus*) shared the central haplotype with ten other sequences; the four other haplotypes were each represented by a single sequence.

### 3.3 Morphological analysis

Based on our morphological analysis, species of the genus *Channichthys* can be divided into two groups according to the structure of the gill apparatus (Fig. 5): (1) that with “double-rowed gill rakers”, having two rows of gill rakers on the gill arches (*sp.br.*) on the outer side of the ceratobranchiale (*sp.br.a*) and on the inner side of the hypobranchiale (*sp.br.b*); and (2) those with “single-rowed gill rakers”, having only one row of gill rakers on the gill arches on the outer side of the ceratobranchiale (*sp.br.a*), but no gill rakers the inner side of the hypobranchiale (*sp.br.b*). Only a single species belongs to the “double-rowed gill rakers” group (*C. panticapaei*), while the “single-rowed gill rakers” group includes three species (*C. rhinoceratus*, *C. rugosus* and *C. velifer*). Thus, these three species are characterized by the first branchial arch bearing only one row of gill rakers on outer side of ceratobranchiale, while rakers are usually absent on the inner side of arch (rarely, single rakers can be found in the angle of the arch).

**Figure 5.**
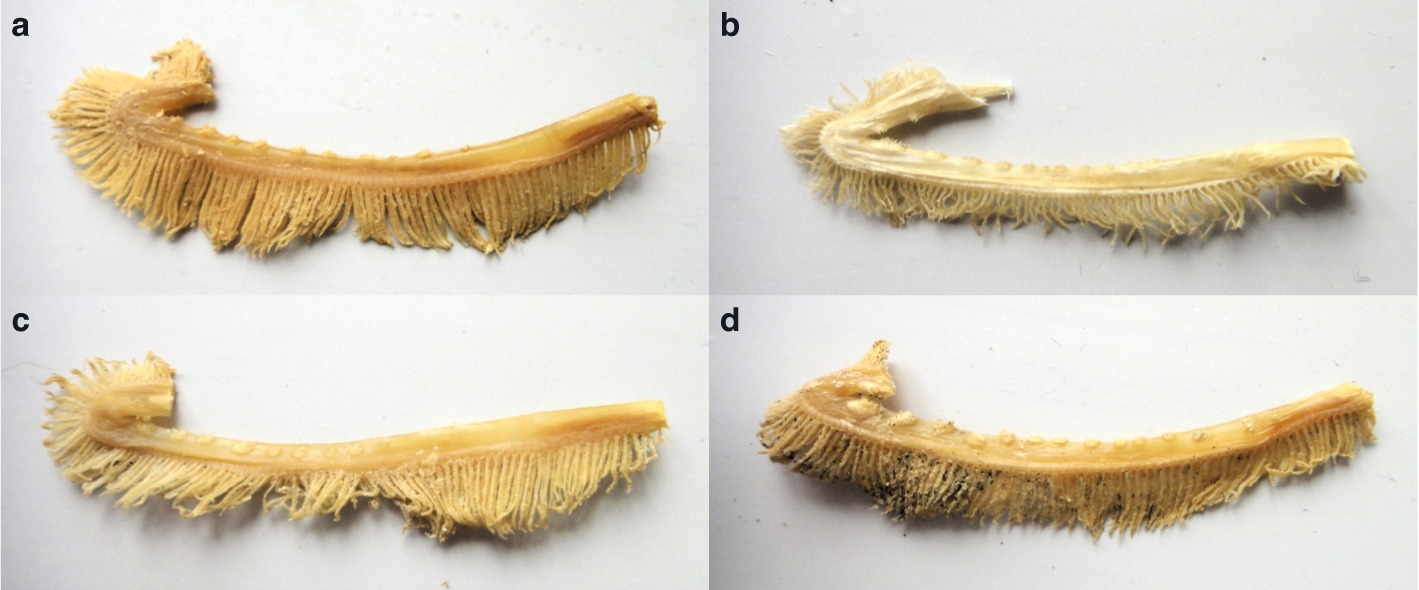
The first gill arches of specimens representing the four recognized species of the genus *Channichthys*. Gill arches are double-rowed in *C. panticapaei* (**d**), but single-rowed in the other three species (**a–c**).

Of all other morphological characteristics (Tables 5 and 6), it is primarily the meristic counts (Table 6) that are of importance for diagnosis, such as: the number of rays in the first (*D1*) and second (*D2*) dorsal fins, in the pectoral fins (*P*), and in the anal fin (*A*); the number of gill rakers on the first gill arches (*sp.br.*), the number of scales in the dorsal (*Dll*) and medial (*Mll*) lateral lines. Additionally, head measurements are important: head length (*c*), head height through the middle of the eye (*ho*), snout length (*ao*), longitudinal diameter orbit of the eye (*o*), interorbital space (*io*), length of the upper (*lmx*) and lower (*lmd*) jaws (Table 5). Finally, certain indices (Table 6) and the external coloration of the body (Fig. 1) can be informative.

**Table 5.** Absolute and relative measurements for the four focal *Channichthys* specimens.

| Character | ZIN 56638<br>(*C. rhinocerus*) | ZIN 56294<br>(*C. rugosus*) | ZIN 56275<br>(*C. velifer*) | ZIN 56538<br>(*C. panticapaei*) |
| --- | --- | --- | --- | --- |
| *TL* (mm) | 420 | 265 | 332 | 363 |
| *SL* (mm) | 380 | 252 | 300 | 320 |
| *ho* (mm) | 43 | 36 | 39 | 35 |
| *hD1* (mm) | 78 | 72 | 73 | 91 |
| *lP<sub>l</sub>* (mm) | 69 | 43 | 54 | 59 |
| *lV<sub>l</sub>* (mm) | 65 | 43 | 55 | 63 |
| *c* (mm) | 142 | 93 | 108 | 116 |
| *o* (mm) | 20 | 15 | 18 | 20 |
| *io* (mm) | 28 | 14 | 19 | 20 |
| *ao* (mm) | 70 | 48 | 52 | 52 |
| *lmx* (mm) | 80 | 51 | 60 | 60 |
| *lmd* (mm) | 97 | 65 | 73 | 77 |
| *ho/SL* | 0.113 | 0.143 | 0.130 | 0.109 |
| *hD1/SL* | 0.205 | 0.286 | 0.243 | 0.284 |
| *lP<sub>l</sub>/SL* | 0.182 | 0.171 | 0.180 | 0.184 |
| *lV<sub>l</sub>/SL* | 0.171 | 0.171 | 0.183 | 0.197 |
| *c/SL* | 0.374 | 0.369 | 0.360 | 0.363 |
| *o/SL* | 0.053 | 0.060 | 0.060 | 0.063 |
| *io/SL* | 0.074 | 0.056 | 0.063 | 0.063 |
| *ao/SL* | 0.184 | 0.190 | 0.173 | 0.163 |
| *lmx/SL* | 0.211 | 0.202 | 0.200 | 0.188 |
| *lmd/SL* | 0.255 | 0.258 | 0.243 | 0.241 |

***TL*** – total body length;
***SL*** – standard body length;
***c*** – head length;
***hD1*** – maximum height of the 1st dorsal fin;
***lP<sub>l</sub>*** and ***lP<sub>d</sub>*** – length of the left and right pectoral fins;
***lV<sub>l</sub>*** and ***lV<sub>d</sub>*** – length of the left and right pelvic fins;
***ho*** – head height through the middle of the eye;
***lmx*** – length of the upper jaw;
***lmd*** – length of the lower jaw;

**Table 6.**
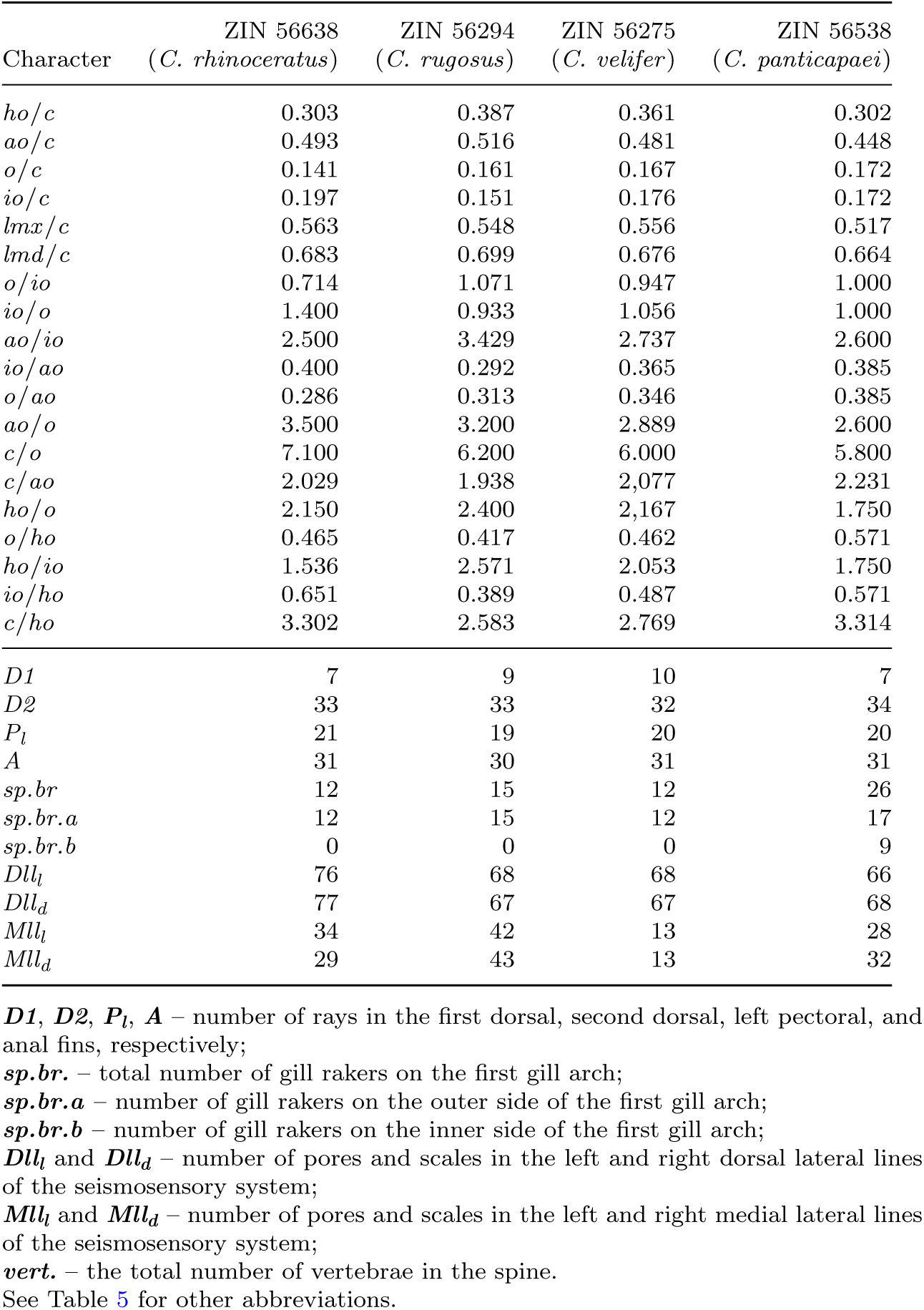
Indices and meristic counts for the four focal *Channichthys* specimens.

*Channichthys rhinoceratus* is characterized by a first dorsal fin (*D1*) with 5–9 (mean 7) rays, a fin membrane that does not reach the tips of the largest rays, a granulation that is poorly or moderately developed, and marbled color.

*Channichthys rugosus* has a first dorsal fin with 7–9 (mean 8) rays, a fin membrane reaching the tips of the largest rays, a granulation that is well developed, and is plain reddish in color.

*Channichthys velifer* has a first dorsal fin with 9–12 (mean 11) rays and a characteristic sail-like shape, a fin membrane that does not reach the tips of the largest rays, poorly developed granulation, and is of lighter spotted color.

As described above, *Channichthys panticapaei* is characterized, in contrast to all other species, by the first branchial arch bearing two rows of gill rakers located on the outer and inner sides of the cerato- and hypobranchiale. The outside rakers count 12–20 and the inside rakers 4–14; the total number of rakers is 18–32. The first dorsal fin has 6–8 (mean 7) rays, the fin membrane does not reach the tips of the largest rays, the granulation is highly developed, and the body is of darker brownish-black color.

Thus, according to morphological characteristics (Tables 5 and 6), four species can be distinguished in the genus *Channichthys*: *C. rhinoceratus*, *C. rugosus*, *C. velifer*, and *C. panticapaei*. A detailed diagnosis of these four species is presented in the Appendix.

## 4 Discussion

Endogenous DNA contents showed large differences among the extracts from the twelve specimens we tested. These differences are surprising given the similar ages of the specimens (50-55 years) and the presumably comparable conditions under which they were fixed and preserved at the Zoological Institute of the Russian Academy of Sciences. These results therefore suggest that DNA degradation may still vary strongly between specimens, likely due to unknown parameters influencing individual degradation levels. Among these could be concentration differences of formaldehyde used for fixation, the fixation duration, the temperature to which specimens were exposed, or variation of ethanol concentration over time in different storage units. Our results, however, show another case, where a DNA concentration below the Qubit High Sensitivity kit detection threshold of 0.05 ng/µl (specimen ZIN 56538) was not impeding successful single-stranded library construction and ultimately obtaining sufficient reads for reconstruction of the mitochondrial genome sequence (Table 2). This has previously been reported (Straube, Preick, et al., 2021) and suggests that samples with undetectable DNA concentration can still be successfully sequenced. In other cases, when shotgun sequencing data indicated low amounts of endogenous DNA (Table 2), target gene capture can be a suitable alternative for future endeavors (e.g., Agne, Naylor, et al., 2022; Agne, Preick, Straube, & Hofreiter, 2022; Straube, Preick, et al., 2021).

Our molecular analyses revealed high similarity, with no more than 34 substitutions in pairwise comparisons, among the mitochondrial genomes of specimens from three of the four recognized species within the genus *Channichthys*: Unicorn Icefish (*C. rhinoceratus*), Red Icefish (*C. rugosus*), and Sailfish Pike (*C. velifer*). Only the specimen representing the Charcoal Icefish (*C. panticapaei*) had a more divergent mitochondrial genome, differing from its congeners at 110–114 sites. These results extend those obtained by Muschick et al. (2022), which had already indicated great similarity between the mitochondrial genomes of Unicorn Icefish (*C. rhinoceratus*) and Red Icefish (*C. rugosus*) specimens.

The lack of substantial mitochondrial variation among three of the four species seems to contrast with their notable morphological differences, in fin ray counts, gill arch structure, size proportions, and coloration, among other traits (see Appendix). We see three possible explanations for this discrepancy. First, the genus *Channichthys* might not include as many species as recognized by recent studies (Nikolaeva, 2020, 2024), even after the synonymization of five formerly recognized taxa (Nikolaeva, 2020; Shandikov, 1995b). This possibility was suggested by Muschick et al. (2022), following their analysis of the mitochondrial genome of *C. rugosus*, and it would mirror conclusions made for another formerly species-rich notothenioid genus, *Pogonophryne* (Parker, Dornburg, Struthers, Jones, & Near, 2022). In Muschick et al. (2022), comparisons of mitochondrial genome sequences revealed that specimens within notothenioid species generally had fewer than 100 genetic differences, whereas between-species comparisons typically exceeded 400 differences. However, the comparisons of Muschick et al. (2022) did not include the closely related, but undoubtedly distinct species pair *Champsocephalus gunnari* and *C. esox* (Rivera-Colón et al., 2023), as no mitochondrial genome was available for the latter at the time of analysis. The now available mitochondrial genome sequences of the two species (NCBI accessions NC 018340 and NC 063099) are separated by 167 substitutions, a number that is far below the over 400 substitutions found in other cross-species comparisons. On the other hand, this number is still not comparable to the 13–34 substitutions separating the mitochondrial genomes of the here analyzed Unicorn Icefish (*C. rhinoceratus*), Red Icefish (*C. rugosus*), and Sailfish Pike (*C. velifer*) specimens. If these three taxa should in fact be part of one and the same species, this species – which, based on priority, would be *C. rhinoceratus* (Richardson, 1844) – would exhibit substantial morphological variation. This variation could perhaps in part be due to phenotypic plasticity, e.g., resulting from life at different depths in the water column influencing food composition or parasite exposure. Such factors are known to affect body shapes (Hetzel & Forsythe, 2023), but are unlikely to influence meristic characters. Better knowledge of the biology of the three taxa, including their (overlapping or not) depth distributions, would be required to assess the extent of phenotypic plasticity in *Channichthys*. Alternatively, large-effect genetic features such as inversion-linked supergenes can produce distinct phenotypes within species (Jeong et al., 2022; Lamichhaney et al., 2016). Such features could be identified with population-level nuclear genomic data, but these data are so far unavailable for the genus *Channichthys*. The most relevant source of information to evaluate *Channichthys* species would be experimental data on potential reproductive barriers; however, the remoteness of their habitat, the difficulty of raising them in captivity, and their late age at sexual maturity render such experiments hard to conclusively perform (Desvignes et al., 2019, 2024). In addition, a few studies have revealed the potential for reproductive isolation between species to be incomplete (Desvignes et al., 2019; Marino et al., 2013; Schiavon et al., 2021) and hybridization barriers to be permissive and potentially driven more by differing reproduction seasons and nesting behavior instead of intrinsic gametic incompatibility (Desvignes et al., 2019).

Second, the three taxa Unicorn Icefish (*C. rhinoceratus*), Red Icefish (*C. rugosus*), and Sailfish Pike (*C. velifer*) might all be distinct and valid species, even though their separation is not reflected in their mitochondrial genomes. Mitochondrial genomes are known to cross species boundaries through hybridization, as long as reproductive barriers remain incomplete (Sloan, Havird, & Sharbrough, 2017). This introgression can lead to mitochondrial capture, the complete replacement of the mitochondrial lineage of one species by that of a closely related one (Willis, Farias, & Ortíı, 2013). Thus, two species that are reproductively largely isolated could have little or no mitochondrial divergence even when nuclear genomes are clearly separated. Again, nuclear genomic data or experimental data on reproductive barriers would be required to evaluate this possibility.

Third, the three *Channichthys* taxa might be valid species that have radiated extremely recently on the Kerguelen-Heard Plateau, in which case both mitochondrial and nuclear divergence would be minimal. Indeed, all four *Channichthys* species inhabit the same geographic area and seemingly the same bathymetric range on the Kerguelen-Heard Plateau, which suggests that the four *Channichthys* species would have diversified in sympatry. This would contrast most other examples of recent divergence in Antarctic notothenioids, that are usually considered to occur in allopatry or parapatry (Corso et al., 2024; Desvignes, Postlethwait, & Konstantinidis, 2020; Dornburg, Federman, Eytan, & Near, 2016; La Mesa, Riginella, & Jones, 2017). We know from other cases of adaptive radiation that large numbers of recognizable species (even when not completely reproductively isolated) can be generated over extremely short time scales, such as the Lake Victoria cichlid fish radiation that produced hundreds of species in no more than 15,000 years (Meier et al., 2023; Ngoepe et al., 2023). However, even this, perhaps most extreme, example of rapid species diversification, resulted in greater mitochondrial variation than present among the three *Channichthys* specimens. While few complete mitochondrial genomes are available for the Lake Victoria cichlid radiation, focusing on the NADH dehydrogenase subunit 2 gene (*mt-nd2*), a marker that has been sequenced for many cichlid species (Wagner, Harmon, & Seehausen, 2012), allows a comparison. At this marker, the three *Channichthys* specimens for the Unicorn Icefish (*C. rhinoceratus*), Red Icefish (*C. rugosus*), and Sailfish Pike (*C. velifer*) differ by no more than 1–2 substitutions. In contrast, representatives of the Lake Victoria cichlid radiation, like *Haplochromis sauvagei*, *Paralabidochromis* sp. ‘rock kribensis’, and *Pundamilia pundamilia*, differ by 4–8 substitutions in their *mt-nd2* sequences (NCBI accessions KJ955418, JQ950392, and KY366728). Thus, unless the radiation of *Channichthys* species would be even more recent than that of Lake Victoria cichlids, their low mitochondrial divergence can not be explained by this scenario.

Regardless of the explanation of low mitochondrial variation among the Unicorn Icefish (*C. rhinoceratus*), Red Icefish (*C. rugosus*), and Sailfish Pike (*C. velifer*) specimens, much more pronounced divergence was observed for the mitochondrial genome of the analyzed Charcoal Icefish (*C. panticapaei*) specimen. This genome differed at 110–114 sites from those of its congeners, a range comparable to the separation of the *Champsocephalus gunnari* and *C. esox* species pair with 167 substitutions and beyond the diversity usually found within a single notothenioid species (Muschick et al., 2022). In light of this molecular separation, together with the distinct morphological variation exhibited by the Charcoal Icefish (*C. panticapaei*), we consider the support for its species status sufficient to recommend its inclusion in the list of validated icefish and more broadly notothenioid species (Eastman & Eakin, 2021). These insights gained from the analyzed museum specimens highlight the importance of natural history collections in biodiversity studies, especially in rarely accessed areas such as Antarctica (Hilton, Watkins-Colwell, & Huber, 2021; O’Brien et al., 2022).

## Acknowledgments

We thank Louise Maria Lindblom (University Museum of Bergen, Norway) and Kenneth Meland (Department of Biology, University of Bergen, Norway) for support during labwork, Michaela Preick (Potsdam University, Germany) for advice and sequencing of DNA libraries, and Moritz Muschick (University of Bern, Switzerland) for helpful discussions on analyses. All computational analyses were performed on resources provided by Sigma2 – the National Infrastructure for High Performance Computing and Data Storage in Norway.

## Author Contributions

BGA contributed to lab work, performed molecular analyses, and wrote a partial draft of the manuscript. EN contributed tissue samples, performed all morphological analyses, and contributed to the draft manuscript. TD contributed conceptually. NS led the lab work and contributed conceptually. MM contributed to lab work, performed molecular analyses, completed the draft manuscript, and contributed conceptually. EN, TD, NS, and MM contributed to the final version of the manuscript.

## Supplementary Information

Supplementary Information is available for this article and includes Supplementary Figures 1–16.

## Appendix

### Differential morphological diagnosis

***Channichthys* Richardson, 1844.**

*D1* 5–12, *D2* 29–35, *P* 17–23, *A* 28–34, *Dll* 56–88, *Mll* 4–45, *sp.br.* 6–32, *vert.* 54–58. The head with an elongated dorsoventrally flattened snout, slightly larger than half the head length, rounded in front and having at the end a well-developed rostral spine with 4–6 separate apical tubercles. The eye size is relatively small. The outer edges of the frontalia above the eyes are slightly raised. The posterior edge of the maxillare usually reaches a vertical line passing through 1/2 of the diameter of the eye orbit. The upper and lower jaws are approximately equal in length, they are covered with 3–10 rows of small, sharp, bristle-like teeth. The 1st and 2nd dorsal fins are well separated; the inter-dorsal space is often relatively wide; the posterior edge of the fin fold of the last ray of the 1st dorsal fin does not reach the base of the 1st ray of the 2nd dorsal fin. The pectoral fins reach the anus. The pelvic fins are wide, slightly shorter or approximately equal to the length of the pectoral fins; they do not reach the anus or end at the level of the anal fin origin. The caudal fin is slightly rounded. There are two lateral lines (dorsal and medial) on the left and right sides along the body (Nikolaeva, 2024).

***Channichthys rhinoceratus* Richardson, 1844.**

This species (Appendix Figure 1) has only a single row of gill rakers (12 on average) on the gill arches (*sp.br.*) (Appendix Figure 2). The first dorsal fin (*D1*) is low, the average relative height (*hD1* /*SL*) is 19.6%; it carries 7 rays on average, of which 2–3 rays (1st to 3rd) are the longest; the fin membrane does not reach the tips of the longest rays. The inter-dorsal space is wide, the first and second dorsal fins (*D1* and *D2*) are well separated. The eye size (*o*) is on average 16.6% of the head length (*c*) or 33.7% of the snout length (*ao*) on average. The interorbital space (*io*) is wide, flat, 16.4% of the head length (*c*) on average; it is often greater than or equal to the eye diameter (*o*). The snout length (*ao*) is slightly less than or approximately equal to half the head length (*c*), averaging 49.3% *c*. The posterior part of the medial lateral lines (*Mll*) usually has moderately developed bony scales. In general, granulation is poorly or moderately developed. The body coloration is usually dark, with numerous dark spots merging into a marbled pattern (Balushkin & Nikolaeva, 2015; Nikolaeva, 2020, 2024).

Morphological characteristics of specimen ZIN 56638 (*C. rhinoceratus*) are provided in Tables 5 and 6.

***Channichthys rugosus* Regan, 1913.**

In this species (Appendix Figure 3), there is also a single row of gill rakers (12 on average) on the gill arches (*sp.br.*) (Appendix Figure 4). The first dorsal fin (*D1*) is of normal shape, high (the relative height *hD1* /*SL* is 20.6% on average); it includes 8 rays on average, the longest ray is usually the fourth one (from the 2nd to the 5th); the fin membrane of *D1* reaches the tips of the longest rays. The inter-dorsal space is wide, *D1* and *D2* are well separated. The eye diameter (*o*) is on average 15.6% of the head length (*c*) or 32.4% of the snout length (*ao*). The interorbital space (*io*) is concave and narrow, averaging 16% of the head length (*c*), it is smaller than the eye diameter (*o*). The snout length (*ao*) is on average 48.2% of the head length (*c*). There are well-developed bony scales along the entire medial lateral lines (*Mll*), including their posterior end. Skin granulation is well developed. The colour of the body and dorsal fins is uniform, species-specific reddish (Nikolaeva, 2021, 2024).

Morphological characteristics of specimen ZIN 56294 (*C. rugosus*) are provided in Tables 5 and 6.

***Channichthys velifer* Meisner, 1974.**

In this species (Appendix Figure 5), there is also a single row of gill rakers (13 on average) on the gill arches (*sp.br.*) (Appendix Figure 6). The first dorsal fin (*D1*) is unique, species-specific (high sail-shaped), with a relative average height (*hD1* /*SL*) of 22.7%; the average number of rays is 10, which is always more than in all other species. The fin membrane between the *D1* rays is high, reaching the tips of the longest rays, 2–5 rays are the longest (from the 3rd to the 7th, usually the 4th or 5th). The inter-dorsal space is very narrow, unlike in other species; the posterior edge of the fin fold of the last ray of *D1* almost reaches the base of the 1st ray of *D2*, so *D1* and *D2* almost touch each other. The eye diameter (*o*) is on average 16.4% of the head length (*c*) or 33.7% of the snout length (*ao*). The interorbital space (*io*) is flat, wide, averaging 17.5% of the head length (*c*); it is usually larger than the eye diameter (*o*). The snout length (*ao*) is on average 48.6% of the head length (*c*). There are usually no bone scales in the posterior part of the medial lateral lines (*Mll*). In general, granulation is poorly or moderately developed. The colour of the body, including the first dorsal fin, is usually lighter than in other species, with numerous scattered small rounded dark spots (Nikolaeva, 2024; Nikolaeva & Balushkin, 2019).

Morphological characteristics of specimen ZIN 56275 (*C. velifer*) are provided in Tables 5 and 6.

***Channichthys panticapaei* Shandikov, 1995a.**

The main species-specific feature that distinguishes *C. panticapaei* (Appendix Figure 7) from all other species of the genus is the presence of two rows of rakers on the gill arches (*sp.br.*): one on the outer side (*sp.br.a*) with 16 rakers on average, and one on the inner side (*sp.br.b*) with 10 rakers on average; the total average number of rakers is 26 (Appendix Figure 8). The first dorsal fin (*D1*) is high, of normal shape, relative average height (*hD1* /*SL*) is 24.6%; the average number of rays is 7, the first three rays (usually the 2nd or 3rd) are the longest; the fin membrane between the rays *D1* is low, does not reach the tips of the longest rays. The inter-dorsal space is relatively wide, *D1* and *D2* are well separated. The eye size (*o*) is slightly larger than in the other species, averaging 18.3% of the head length (*c*) or averaging 39.3% of the snout length (*ao*). The interorbital space (*io*) is wide, flat, but more often equal to the eye diameter (*o*), averaging 16.9% of the head length (*c*). The snout length (*ao*) is slightly less than that of other species, averaging 46.8% of the head length (*c*). Along the entire medial lateral lines (*Mll*), including their posterior end, there are bone scales developed very strongly. Granulation is generally highly developed. The body colour is very characteristic, it is darker than in other species: brown-black, especially the upper part of the head and body, the back is covered with even darker spots, merging into a marble pattern (Nikolaeva, 2019, 2024).

Morphological characteristics of specimen ZIN 56538 (*C. panticapaei*) are provided in Tables 5 and 6.

---

**Appendix Figure 1.**
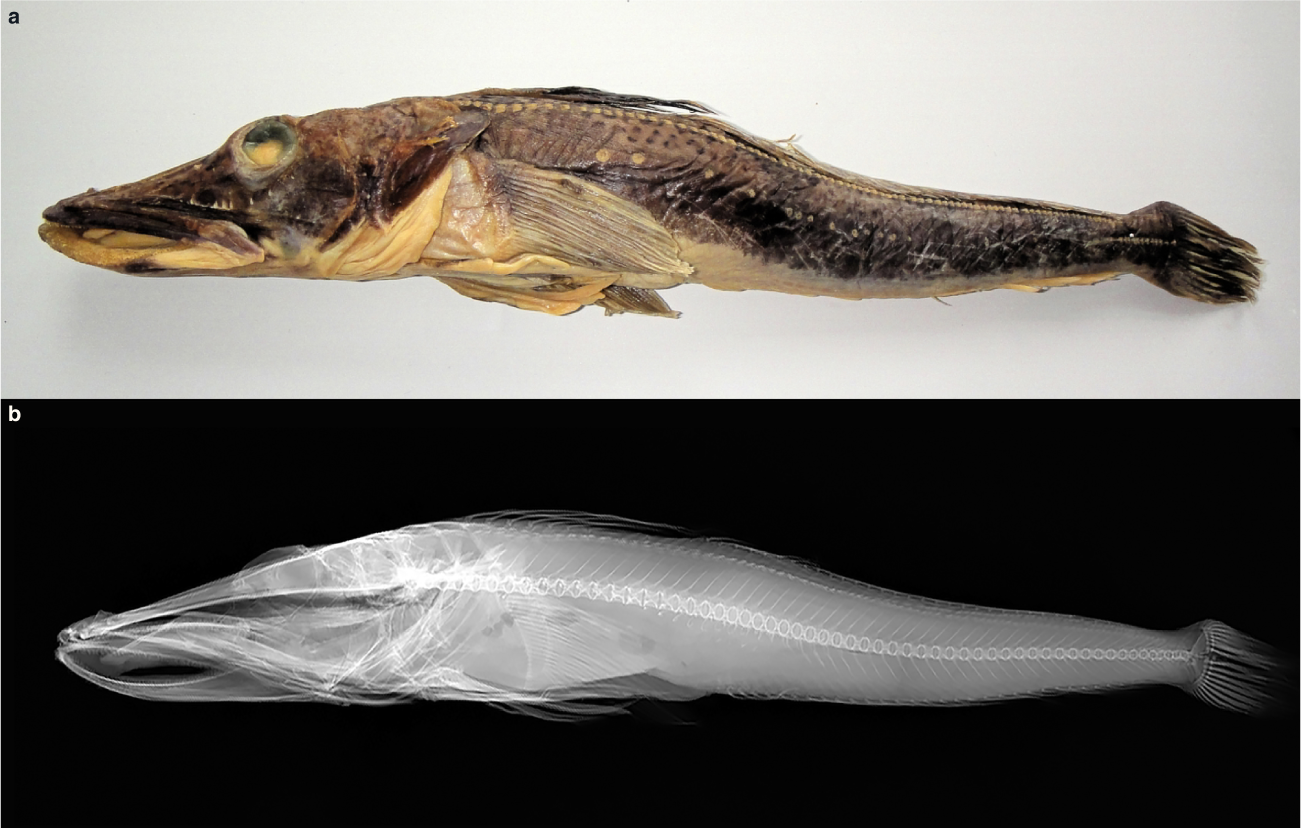
ZIN 56638 (*C. rhinoceratus*). **a** Photograph; **b** X-ray image.

**Appendix Figure 2.**
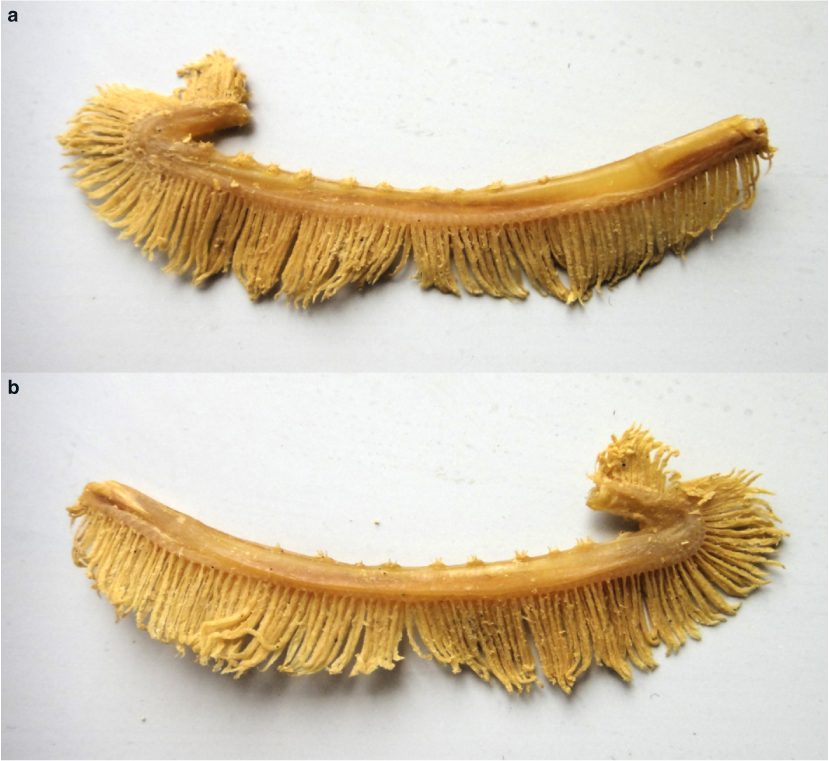
First gill arch of ZIN 56638 (*C. rhinoceratus*). **a** External side; **b** internal side.

**Appendix Figure 3.**
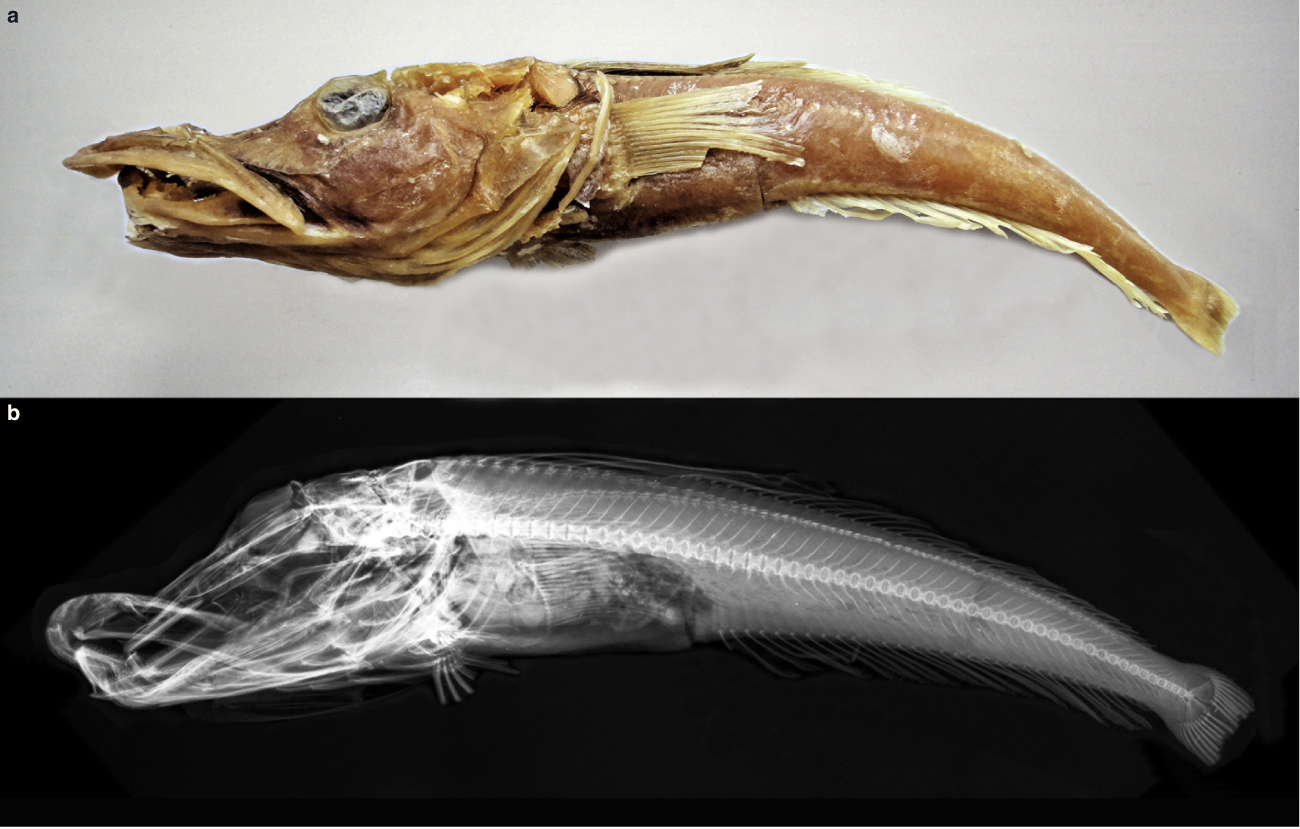
ZIN 56294 (*C. rugosus*). **a** Photograph; **b** X-ray image.

**Appendix Figure 4.**
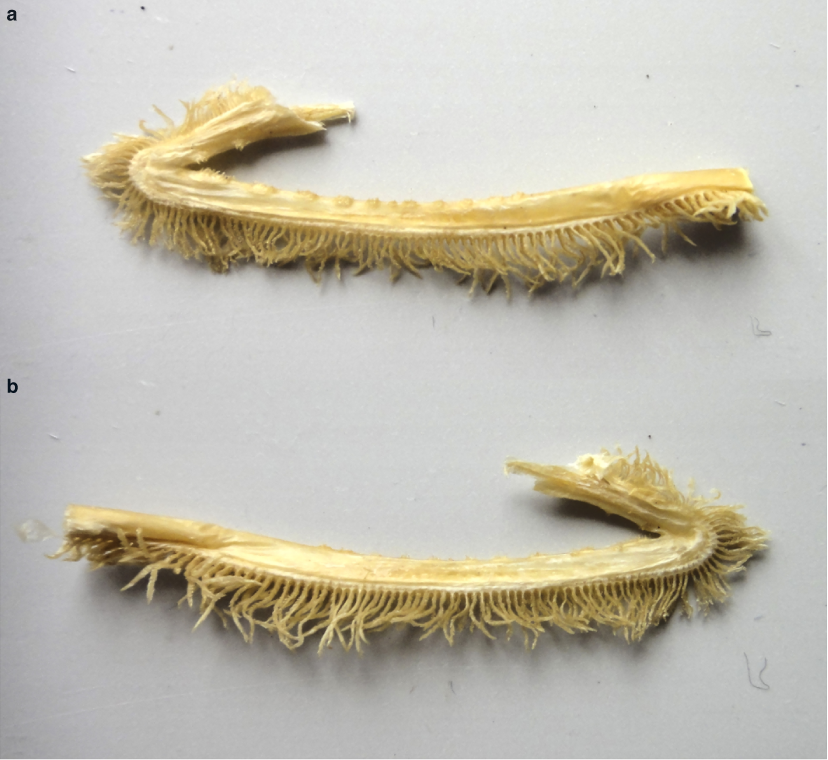
First gill arch of ZIN 56294 (*C. rugosus*). **a** External side; **b** internal side.

**Appendix Figure 5.**
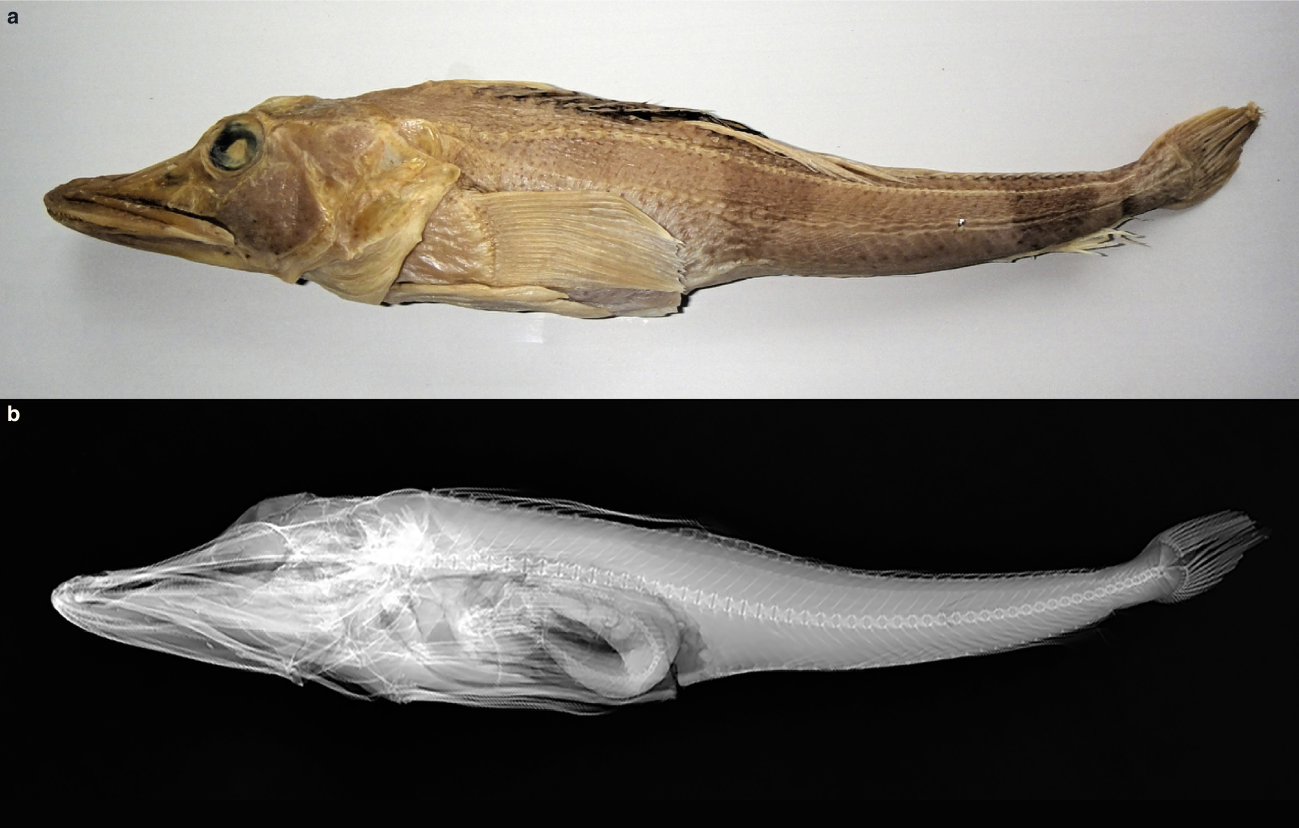
ZIN 56275 (*C. velifer*). **a** Photograph; **b** X-ray image.

**Appendix Figure 6.**
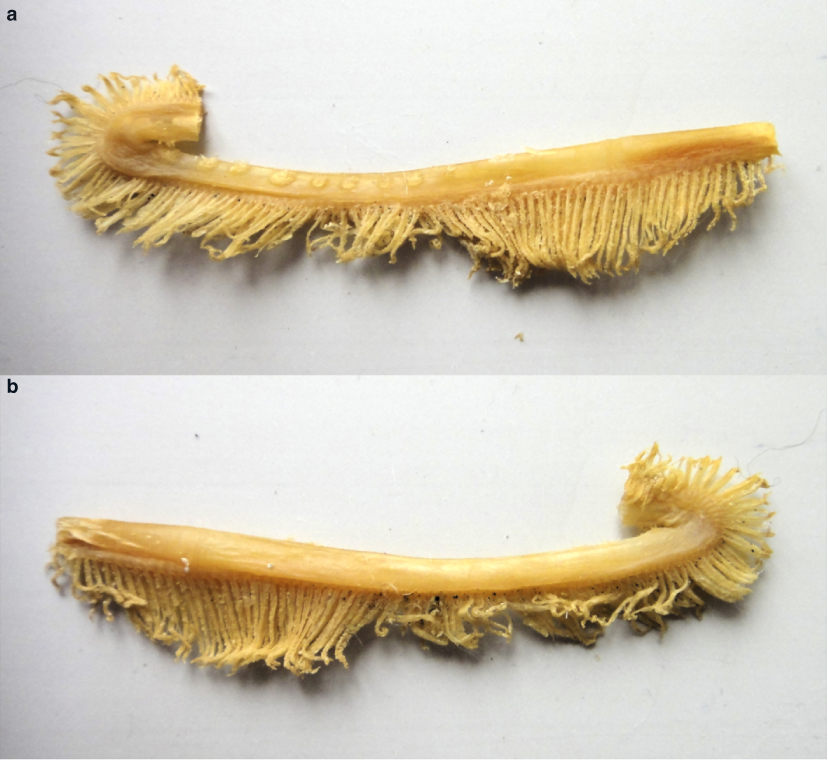
First gill arch of ZIN 56275 (*C. velifer*). **a** External side; **b** internal side.

**Appendix Figure 7.**
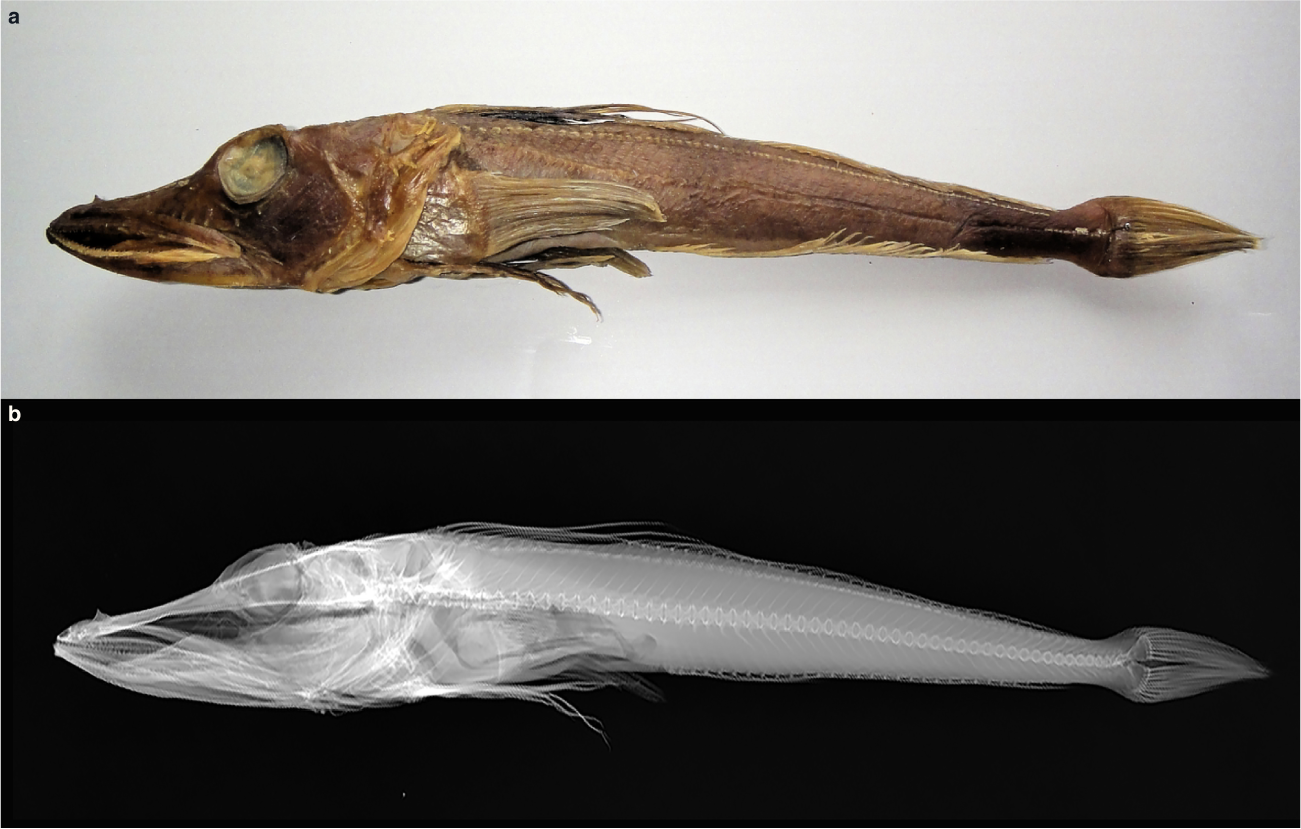
ZIN 56538 (*C. panticapaei*). **a** Photograph; **b** X-ray image.

**Appendix Figure 8.**
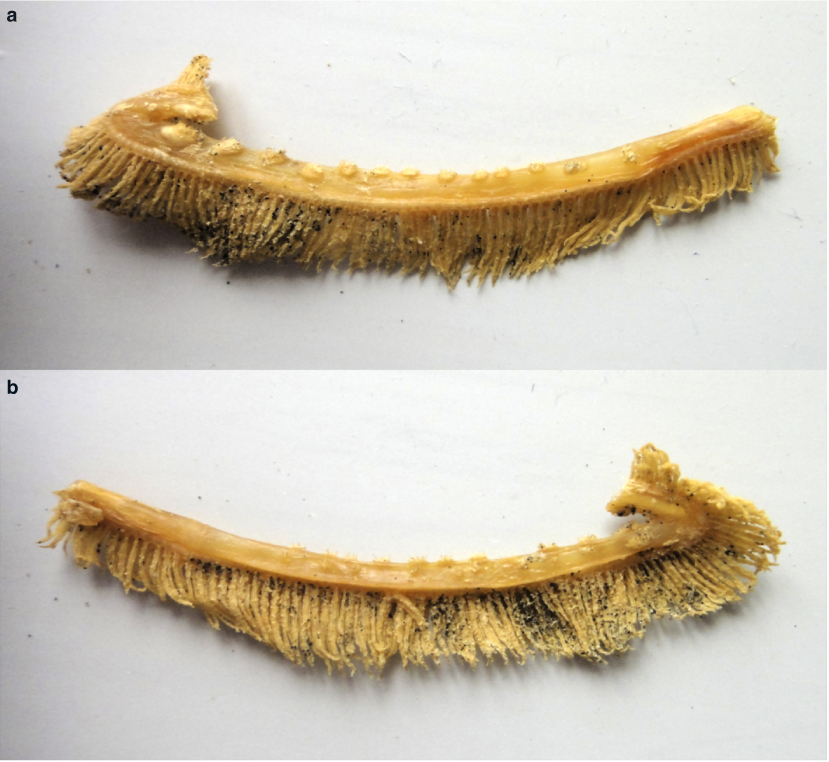
First gill arch of ZIN 56538 (*C. panticapaei*). **a** External side; **b** internal side.

## References

Agne, S., Naylor, G.J.P., Preick, M., Yang, L., Thiel, R., Weigmann, S.,…Straube, N. (2022). Taxonomic identification of two poorly known lantern shark species based on mitochondrial DNA from wet-collection paratypes. Frontiers in Ecology and Evolution, 10, 910009. 10.3389/fevo.2022.910009

Agne, S., Preick, M., Straube, N., Hofreiter, M. (2022). Simultaneous barcode sequencing of diverse museum collection specimens using a mixed RNA bait set. Frontiers in Ecology and Evolution, 10, 909846. 10.3389/fevo.2022.909846

Ahyong, S., Boyko, C., Bernot, J., Brandão, S., Daly, M., De Grave, S.,…Zullini, A. (2024). World Register of Marine Species (WoRMS). WoRMS Editorial Board. (Accessed: 2024-09-08)

Altschul, S.F., Gish, W., Miller, W., Myers, E.W., Lipman, D.J. (1990). Basic local alignment search tool. Journal of Molecular Biology, 215 (3), 403– 410. 10.1016/s0022-2836(05)80360-2

Balushkin, A.V., & Nikolaeva, E.A. (2015). “Dolichobranchiata” mutation in the Antarctic representatives from the families of plunderfishes (Artedidraconidae)and white-blooded (Channichthyidae) fish (Notothenioidei). Journal of Ichthyology, 55 (1), 9–15. 10.1134/S0032945215010014

Balushkin, A.V., & Spodareva, V.V. (2015, November). New species of the toad plunderfish of the “albipinna” group, genus *Pogonophryne* (Artedidraconidae) from the Ross Sea (Antarctica). Journal of Ichthyology, 55 (6), 757–764. 10.1134/S003294521506003X

Basler, N., Xenikoudakis, G., Westbury, M.V., Song, L., Sheng, G., Barlow, A. (2017). Reduction of the contaminant fraction of DNA obtained from an ancient giant panda bone. BMC Research Notes, 10 (1), 754. 10.1186/s13104-017-3061-3

Bista, I., Wood, J.M.D., Desvignes, T., McCarthy, S.A., Matschiner, M., Ning, Z.,…Durbin, R. (2023). Genomics of cold adaptations in the Antarctic notothenioid fish radiation. Nature Communications, 14, 3412. 10.1101/2022.06.08.494096

Buchfink, B., Xie, C., Huson, D.H. (2015). Fast and sensitive protein alignment using DIAMOND. Nature Methods, 12 (1), 59–60. 10.1038/nmeth.3176

Chen, S., Zhou, Y., Chen, Y., Gu, J. (2018). fastp: an ultra-fast all-in-one FASTQ preprocessor. Bioinformatics, 34 (17), i884–i890. Retrieved 2022-10-24, from https://academic.oup.com/bioinformatics/article/34/17/i884/5093234 10.1093/bioinformatics/bty560

Clarke, A., & Johnston, I.A. (1996). Evolution and adaptive radiation of antarctic fishes. Trends in Ecology and Evolution, 11 (5), 212–218. 10.1016/0169-5347(96)10029-X

Colombo, M., Damerau, M., Hanel, R., Salzburger, W., Matschiner, M. (2015). Diversity and disparity through time in the adaptive radiation of Antarctic notothenioid fishes. Journal of Evolutionary Biology, 28 (2), 376–394. 10.1111/jeb.12570

Corso, A.D., Desvignes, T., Mcdowell, J.R., Cheng, C.-H.C., Biesack, E.E., Steinberg, D.K., Hilton, E.J. (2024). *Akarotaxis gouldae*, a new species of Antarctic dragonfish (Notothenioidei: Bathydraconidae) from the western Antarctic Peninsula. Zootaxa, 5501 (2), 265–290. 10.11646/zootaxa.5501.2.3

Dabney, J., Knapp, M., Glocke, I., Gansauge, M.-T., Weihmann, A., Nickel, B.,…Meyer, M. (2013). Complete mitochondrial genome sequence of a Middle Pleistocene cave bear reconstructed from ultrashort DNA fragments. Proceedings of the National Academy of Sciences, 110 (39), 15758–15763. 10.1073/pnas.1314445110

Desvignes, T., Bista, I., Herrera, K., Landes, A., Postlethwait, J.H. (2023). Cold-driven hemoglobin evolution in Antarctic notothenioid fishes prior to hemoglobin gene loss in white-blooded icefishes. Molecular Biology and Evolution, 40 (11), msad236. 10.1093/molbev/msad236

Desvignes, T., Le François, N.R., Goetz, L.C., Smith, S.S., Shusdock, K.A., Parker, S.K.,…Detrich, H.W. (2019). Intergeneric hybrids inform reproductive isolating barriers in the Antarctic icefish radiation. Scientific Reports, 9 (1), 5989. 10.1038/s41598-019-42354-z

Desvignes, T., Le François, N.R., Streeter, M., Grondin, J., Singer, E., Postlethwait, J.H., Detrich, H.W. (2024). Hybridization barriers between the congeneric antarctic notothenioid fish *Notothenia* coriiceps and *Notothenia* rossii. Polar Biology, 47 (2), 163–171. 10.1007/s00300-023-03216-7

Desvignes, T., Postlethwait, J.H., Konstantinidis, P. (2020). Biogeography of the Antarctic dragonfishes Acanthodraco dewitti and Psilodraco breviceps with re-description of Acanthodraco dewitti larvae (Notothenioidei: Bathydraconidae). Polar Biology, 43 (5), 565–572. 10.1007/s00300-020-02661-y

DeVries, A.L., & Wohlschlag, D.E. (1969). Freezing resistance in some Antarctic fishes. Science, 163 (871), 1073–1075. 10.1126/science.163.3871.107

Dornburg, A., Federman, S., Eytan, R.I., Near, T.J. (2016). Cryptic species diversity in sub-Antarctic islands: A case study of *Lepidonotothen*. Molecular Phylogenetics and Evolution, 104, 32–43. 10.1016/j.ympev.2016.07.013

Dornburg, A., Federman, S., Lamb, A.D., Jones, C.D., Near, T.J. (2017). Cradles and museums of Antarctic teleost biodiversity. Nature Ecology and Evolution, 1, 1379–1384. 10.1038/s41559-017-0239-y

Duhamel, G., Gasco, N., & Davaine, P. (Eds.). (2005). Poissons des I^^^les Kerguelen et Crozet. Guide Ŕegional de l’Oćean Austral. Patrimoines Naturels (Vol. 63). Paris: Muśeum National d’Histoire Naturelle.

Eastman, J.T. (1993). Antarctic Fish Biology: Evolution in a Unique Environment. San Diego, CA: Academic Press, Inc.

Eastman, J.T. (2005). The nature of the diversity of Antarctic fishes. Polar Biology, 28 (2), 93–107. 10.1007/s00300-004-0667-4

Eastman, J.T., & Eakin, R.R. (2021). Checklist of the species of notothenioid fishes. Antarctic Science, 33 (3), 273–280. 10.1017/S0954102020000632

Fricke, R., Eschmeyer, W.N., van der Laan, R. (2024). Eschmeyer’s Catalog of Fishes. California Academy of Sciences. (Accessed: 2024–09-08)

Froese, R., & Pauly, D. (2024). Fishbase. World Wide Web electronic publication. (Accessed: 2024-09-08)

Fulton, T.L., & Shapiro, B. (2019). Setting up an ancient DNA laboratory. B. Shapiro, A. Barlow, P.D. Heintzman, M. Hofreiter, J.L.A. Paijmans, & A.E.R. Soares (Eds.), Ancient DNA (Vol. 1963, pp. 1–13). New York, NY: Springer New York. 10.1007/978-1-4939-9176-1 1

Gansauge, M.-T., Aximu-Petri, A., Nagel, S., Meyer, M. (2020). Manual and automated preparation of single-stranded DNA libraries for the sequencing of DNA from ancient biological remains and other sources of highly degraded DNA. Nature Protocols, 15 (8), 2279–2300. 10.1038/s41596-020-0338-0

Gansauge, M.-T., Gerber, T., Glocke, I., Korlević, P., Lippik, L., Nagel, S.,…Meyer, M. (2017). Single-stranded DNA library preparation from highly degraded DNA using *T4* DNA ligase. Nucleic Acids Research, gkx033. 10.1093/nar/gkx033

Grant, P., & Grant, B. (2008). How and why species multiply: the radiation of Darwin’s finches. Princeton, New Jersey, USA: Princeton University Press.

Hetzel, C., & Forsythe, P. (2023). Phenotypic plasticity of a generalist fish species resident to lotic environments: Insights from the Great Lakes region. Ecology and Evolution, 13 (11), e10715. 10.1002/ece3.10715

Hilton, E.J., Watkins-Colwell, G.J., Huber, S.K. (2021). The expanding role of natural history collections. Ichthyology & Herpetology, 109 (2). 10.1643/t2020018

Huson, D.H., Beier, S., Flade, I., Górska, A., El-Hadidi, M., Mitra, S.,…Tappu, R. (2016). MEGAN Community Edition - interactive exploration and analysis of large-scale microbiome sequencing data. PLOS Computational Biology, 12 (6), e1004957. 10.1371/journal.pcbi.1004957

Jeong, H., Baran, N.M., Sun, D., Chatterjee, P., Layman, T.S., Balakrishnan, C.N.,…Yi, S.V. (2022). Dynamic molecular evolution of a supergene with suppressed recombination in white-throated sparrows. eLife, 11. 10.7554/eLife.79387

Jónsson, H., Ginolhac, A., Schubert, M., Johnson, P.L.F., Orlando, L. (2013). mapDamage2.0: fast approximate Bayesian estimates of ancient DNA damage parameters. Bioinformatics, 29 (13), 1682 – 1684. 10.1093/bioinformatics/btt193

Katoh, K., & Standley, D.M. (2013). MAFFT multiple sequence alignment software version 7: improvements in performance and usability. Molecular Biology and Evolution, 30 (4), 772–780. 10.1093/molbev/mst010

La Mesa, M., Riginella, E., Jones, C.D. (2017). Early life history traits and geographical distribution of *Parachaenichthys charcoti*. Antarctic Science, 29 (5), 410–416. 10.1017/S0954102017000189

Lamichhaney, S., Fan, G., Widemo, F., Gunnarsson, U., Thalmann, D.S., Hoeppner, M.P.,…Andersson, L. (2016). Structural genomic changes underlie alternative reproductive strategies in the ruff (Philomachus pugnax). Nature Genetics, 48 (1), 84–88. 10.1038/ng.3430

Lerner, H., Meyer, M., James, H., Hofreiter, M., Fleischer, R. (2011). Multilocus resolution of phylogeny and timescale in the extant adaptive radiation of Hawaiian honeycreepers. Current Biology, 21 (21), 1838–1844. 10.1016/j.cub.2011.09.039

Li, H. (2011). A statistical framework for SNP calling, mutation discovery, association mapping and population genetical parameter estimation from sequencing data. Bioinformatics, 27 (21), 2987–2993. 10.1093/bioinformatics/btr509

Li, H., & Durbin, R. (2010). Fast and accurate long-read alignment with Burrows-Wheeler transform. Bioinformatics, 26 (5), 589 – 595. 10.1093/bioinformatics/btp698

Losos, J. (2009). Lizards in an Evolutionary Tree. Berkeley, California, USA: University of California Press.

Marino, I.A.M., Benazzo, A., Agostini, C., Mezzavilla, M., Hoban, S.M., Patarnello, T.,…Bertorelle, G. (2013). Evidence for past and present hybridization in three Antarctic icefish species provides new perspectives on an evolutionary radiation. Molecular Ecology, 22 (20), 5148–5161. 10.1111/mec.12458

Matschiner, M. (2016). Fitchi: haplotype genealogy graphs based on the Fitch algorithm. Bioinformatics, 32 (8), 1250–1252. 10.1093/bioinformatics/btv717

Matschiner, M., Colombo, M., Damerau, M., Ceballos, S., Hanel, R., Salzburger, W. (2015). The adaptive radiation of notothenioid fishes in the waters of Antarctica. Extremophile Fishes: Ecology, Evolution, and Physiology of Teleosts in Extreme Environments (Vol. 36). Springer International Publishing. 10.1111/j.1365-294X.2006.03105.x

McKenna, A., Hanna, M., Banks, E., Sivachenko, A., Cibulskis, K., Kernytsky, A.,…DePristo, M.A. (2010). The Genome Analysis Toolkit: A MapReduce framework for analyzing next-generation DNA sequencing data. Genome Research, 20 (9), 1297–1303. 10.1101/gr.107524.110

Meier, J.I., McGee, M.D., Marques, D.A., Mwaiko, S., Kishe, M., Wandera, S.,…Seehausen, O. (2023). Cycles of fusion and fission enabled rapid parallel adaptive radiations in African cichlids. Science, 381 (6665), eade2833. 10.1126/science.ade2833

Meisner, E.E. (1974). New species of the icefishes from the Southern Ocean. Vestnik Zoologii, 6, 50–55.

Minh, B.Q., Nguyen, M.A.T., von Haeseler, A. (2013). Ultrafast approximation for phylogenetic bootstrap. Molecular Biology and Evolution, 30 (5), 1188–1195. 10.1093/oxfordjournals.molbev.a025811

Minh, B.Q., Schmidt, H.A., Chernomor, O., Schrempf, D., Woodhams, M.D., Von Haeseler, A., Lanfear, R. (2020). IQ-TREE 2: New models and efficient methods for phylogenetic inference in the genomic era. Molecular Biology and Evolution, 37 (5), 1530–1534. 10.1093/molbev/msaa015

Muschick, M., Nikolaeva, E., Rüber, L., Matschiner, M. (2022). The mitochondrial genome of the red icefish (*Channichthys rugosus*) casts doubt on its species status. Polar Biology, 45, 1541–1552. 10.1007/s00300-022-03083-8

Near, T.J., Dornburg, A., Harrington, R.C., Oliveira, C., Pietsch, T.W., Thacker, C.E.,…Beaulieu, J.M. (2015). Identification of the notothenioid sister lineage illuminates the biogeographic history of an Antarctic adaptive radiation. BMC Evolutionary Biology, 15, 109. 10.1186/s12862-015-0362-9

Near, T.J., Dornburg, A., Kuhn, K.L., Eastman, J.T., Pennington, J.N., Patarnello, T.,…Jones, C.D. (2012). Ancient climate change, antifreeze, and the evolutionary diversification of Antarctic fishes. Proceedings of the National Academy of Sciences USA, 109 (9), 3434–3439. 10.1073/pnas.1115169109

Near, T.J., MacGuigan, D.J., Parker, E., Struthers, C.D., Jones, C.D., Dornburg, A. (2018). Phylogenetic analysis of Antarctic notothenioids illuminates the utility of RADseq for resolving Cenozoic adaptive radiations. Molecular Phylogenetics and Evolution, 129, 268–279. 10.1016/j.ympev.2018.09.001

Ngoepe, N., Muschick, M., Kishe, M.A., Mwaiko, S., Temoltzin-Loranca, Y., King, L.,…Seehausen, O. (2023). A continuous fish fossil record reveals key insights into adaptive radiation. Nature. 10.1038/s41586-023-06603-6

Nikolaeva, E.A. (2016). Systematics of the Kerguelen icefishes of the genus Channichthys Richardson, 1844 (fam. Channichthyidae). O.V. Zaitseva & A.A. Petrov (Eds.), Materials of the 3th All-Russian Conference: Modern Problems in Evolutionary Morphology of Animals (pp. 86–87). Saint Petersburg, Russia: Zoological Institute RAS, Saint Petersburg.

Nikolaeva, E.A. (2017). Taxonomic revision of the Antarctic icefishes of the genus Channichthys Richardson, 1844 (fam. Channichthyidae). S.Y. Sinev & M.K. Stanyukovich (Eds.), Materialy yubileynoy otchetnoy Nauchnoy Sessii, Posvyashchennoy 185-letiyu Zoologicheskogo Instituta RAN (pp. 134–137). Saint Petersburg, Russia: Zoological Institute RAS, Saint Petersburg.

Nikolaeva, E.A. (2019). A review of the icefish species from the genus Channichthys Richardson, 1844 (Channichthyidae) with double-rowed gill rakers. Proceedings of the Zoological Institute RAS, 323 (4), 558–567. 10.31610/trudyzin/2019.323.4.558

Nikolaeva, E.A. (2020). Redescription of the unicorn icefish *Channichthys rhinoceratus* Richardson (Notothenioidei: Channichthyidae) with synonymization of three similar species. Proceedings of the Zoological Institute RAS, 324 (4), 485–496. 10.31610/trudyzin/2020.324.4.485

Nikolaeva, E.A. (2021). On the taxonomic status of the red icefish *Channichthys rugosus* (Notothenioidei: Channichthyidae) from the Kerguelen Islands (South Ocean). Proceedings of the Zoological Institute RAS, 325, 485–494. 10.31610/trudyzin/2021.325.4.485

Nikolaeva, E.A. (2024). Overview of Antarctic icefish species of the genus *Channichthys* Richardson, 1844 (Notothenioidei: Channichthyidae) based on morphological studies. Preprint.

Nikolaeva, E.A., & Balushkin, A.V. (2019). Morphological characteristics of sailfish pike *Channichthys velifer* (Channichthyidae) from the Kerguelen Islands (Southern Ocean). Journal of Ichthyology, 59 (6), 834–842. 10.1134/S0032945219060079

O’Brien, K.M., Crockett, E.L., Adams, B.J., Amsler, C.D., Appiah-Madson, H.J., Collins, A., Watkins-Colwell, G.J. (2022). The time is right for an Antarctic biorepository network. Proceedings of the National Academy of Sciences, 119 (50), e2212800119. 10.1073/pnas.2212800119

Paijmans, J.L.A., Baleka, S., Henneberger, K., Taron, U.H., Trinks, A., Westbury, M.V., Barlow, A. (2017). Sequencing single-stranded libraries on the Illumina NextSeq 500platform. arXiv:1711.11004 [q-bio]. (arXiv: 1711.11004)

Papetti, C., Babbucci, M., Dettaï, A., Basso, A., Lucassen, M., Harms, L.,…Negrisolo, E. (2021). Not frozen in the ice: large and dynamic rearrangements in the mitochondrial genomes of the Antarctic fish. Genome Biology and Evolution, 13 (3), 1–21. 10.1093/gbe/evab017

Parker, E., Dornburg, A., Struthers, C.D., Jones, C.D., Near, T.J. (2022). Phylogenomic species delimitation dramatically reduces species diversity in an Antarctic adaptive radiation. Systematic Biology, 71 (1), 58–77. 10.1093/sysbio/syab057

Rankin, J.C., & Tuurala, H. (1998). Gills of Antarctic fish. Comparative Biochemistry and Physiology, 119A, 149–163.

Regan, C.T. (1913). II. – The Antarctic fishes of the Scottish National Antarctic expedition. Transactions of the Royal Society of Edinburgh: Earth Sciences, 49 (2), 229–292.

Richardson, J. (1844, June). LII.— Descriptions of a new Genus of Gobioid Fish. Annals and Magazine of Natural History, 13 (86), 461–462. 10.1080/03745484409442631

Rivera-Colón, A.G., Rayamajhi, N., Minhas, B.F., Madrigal, G., Bilyk, K.T., Yoon, V.,…Catchen, J.M. (2023). Genomics of secondarily temperate adaptation in the only non-Antarctic icefish. Molecular Biology and Evolution, 40 (3), msad029. 10.1093/molbev/msad029

Robinson, J.T., Thorvaldsdóttir, H., Winckler, W., Guttman, M., Lander, E.S., Getz, G., Mesirov, J.P. (2011). Integrative genomics viewer. Nature Biotechnology, 29, 24–26. 10.1038/nbt0111-24

Rohland, N., Glocke, I., Aximu-Petri, A., Meyer, M. (2018). Extraction of highly degraded DNA from ancient bones, teeth and sediments for high-throughput sequencing. Nature Protocols, 13 (11), 2447–2461. 10.1038/s41596-018-0050-5

Rohland, N., & Hofreiter, M. (2007a). Ancient DNA extraction from bones and teeth. Nature Protocols, 2 (7), 1756–1762. 10.1038/nprot.2007.247

Rohland, N., & Hofreiter, M. (2007b). Comparison and optimization of ancient DNA extraction. BioTechniques, 42 (3), 343–352. 10.2144/000112383

Rohland, N., Siedel, H., Hofreiter, M. (2010). A rapid column-based ancient DNA extraction method for increased sample throughput. Molecular Ecology Resources, 10 (4), 677–683. 10.1111/j.1755-0998.2009.02824.x

Ronco, F., Matschiner, M., Böhne, A., Boila, A., Büscher, H.H., El Taher, A.,…Salzburger, W. (2021). Drivers and dynamics of a massive adaptive radiation in cichlid fishes. Nature, 589, 76–81. 10.1038/s41586-020-2930-4

Ruud, J.T. (1954). Vertebrates without erythrocytes and blood pigment. Nature, 173 (4410), 848–850. 10.1038/173848a0

Sambrook, J., & Russell, D.W. (2001). Molecular cloning: a laboratory manual (3rd ed ed.). Cold Spring Harbor, N.Y: Cold Spring Harbor Laboratory Press.

Schiavon, L., Dulière, V., La Mesa, M., Marino, I.A.M., Codogno, G., Boscari, E.,…Papetti, C. (2021). Species distribution, hybridization and connectivity in the genus *Chionodraco*: Unveiling unknown icefish diversity in antarctica. Diversity and Distributions, 27 (5), 766–783. 10.1111/ddi.13249

Schluter, D. (2000). The Ecology of Adaptive Radiation. New York, NY, USA: Oxford University Press.

Schubert, M., Ermini, L., Der Sarkissian, C., Jónsson, H., Ginolhac, A., Schaefer, R.,…Orlando, L. (2014). Characterization of ancient and modern genomes by SNP detection and phylogenomic and metagenomic analysis using PALEOMIX. Nature Protocols, 9 (5), 1056–1082. 10.1038/nprot.2014.063

Schubert, M., Lindgreen, S., Orlando, L. (2016). AdapterRemoval v2: rapid adapter trimming, identification, and read merging. BMC Research Notes, 9 (1), 88. 10.1186/s13104-016-1900-2

Shandikov, G.A. (1995a). A new species of icefish, *Channichthys panticapei* sp. n. (Channichthyidae, Notothenioidei) from the Kerguelen Island (Antarctica). *Proceedings of South Research Institute of Marine Fishery and Oceanography (YugNIRO)*, Special Issue, 1, 1–10.

Shandikov, G.A. (1995b). To the question about the composition of icefish species of the genus *Channichthys* in the Kerguelen Islands area with description of three new species. Proceedings of South Research Institute of Marine Fishery and Oceanography (YugNIRO), Special Issue, 2, 1–18.

Shandikov, G.A. (2008). *Channichthys mithridatis*, a new species of icefishes (Perciformes: Notothenioidei: Channichthyidae) from the Kerguelen Islands (East Antarctica), with comments on the taxonomic status of *Channichthys normani*. Visnyk Charkivs’koho Universytetu Imeni V. N. Karazina, Ser. Biologija, Charkiv, 14 (917), 123–131.

Shandikov, G.A. (2011). *Channichthys richardsoni* sp. n., a new Antarctic icefish (Perciformes: Notothenioidei: Channichthyidae) from the Kerguelen Islands area, Indian sector of the Southern Ocean. Journal of V. N. Karazin Kharkiv National University, Series: Biology (14), 125–134.

Sidell, B.D., & O’Brien, K. (2006). When bad things happen to good fish: the loss of hemoglobin and myoglobin expression in Antarctic icefishes. Journal of Experimental Biology, 209 (10), 1791–1802. 10.1242/jeb.02091

Simpson, G.G. (1953). The Major Features of Evolution. New York, USA: Columbia University Press.

Sloan, D.B., Havird, J.C., Sharbrough, J. (2017). The on-again, off-again relationship between mitochondrial genomes and species boundaries. Molecular Ecology, 26 (8), 2212–2236. 10.1111/mec.13959

Smith, P.J., Steinke, D., Dettai, A., McMillan, P., Welsford, D., Stewart, A., Ward, R.D. (2012, September). DNA barcodes and species identifications in Ross Sea and Southern Ocean fishes. Polar Biology, 35 (9), 1297–1310. 10.1007/s00300-012-1173-8

Straube, N., Lyra, M.L., Paijmans, J.L.A., Preick, M., Basler, N., Penner, J.,…Hofreiter, M. (2021). Successful application of ancient DNA extraction and library construction protocols to museum wet collection specimens. Molecular Ecology Resources, 21 (7), 2299–2315. 10.1111/1755-0998.13433

Straube, N., Preick, M., Naylor, G.J.P., Hofreiter, M. (2021). Mitochondrial DNA sequencing of a wet-collection syntype demonstrates the importance of type material as genetic resource for lantern shark taxonomy (Chondrichthyes: Etmopteridae). Royal Society Open Science, 8 (9), 210474. 10.1098/rsos.210474

Tuta, B., Acierno, R., Agnisola, C. (1991, June). Mechanical performance of the isolated and perfused heart of the haemoglobinless Antarctic icefish *Chionodraco hamatus* (Lönnberg): effects of loading conditions and temperature. Philosophical Transactions of the Royal Society of London. Series B: Biological Sciences, 332 (1264), 191–198. 10.1098/rstb.1991.0049

Vences, M., Patmanidis, S., Schmidt, J.-C., Matschiner, M., Miralles, A., Renner, S.S. (2024). Hapsolutely: a user-friendly tool integrating haplotype phasing, network construction, and haploweb calculation. Bioinformatics Advances, 4 (1), vbae083. 10.1093/bioadv/vbae083

Wagner, C.E., Harmon, L.J., Seehausen, O. (2012). Ecological opportunity and sexual selection together predict adaptive radiation. Nature, 487 (7407), 366–369. 10.1038/nature11144

Willis, S.C., Farias, I.P., Ortí, G. (2013). Testing mitochondrial capture and deep coalescence in Amazonian cichlid fishes (Cichlidae *Cichla*). Evolution, 68 (1), 256–268. 10.1111/evo.12230

Zhu, T., Sato, Y., Sado, T., Miya, M., Iwasaki, W. (2023, March). MitoFish, MitoAnnotator, and MiFish pipeline: Updates in 10 years. Molecular Biology and Evolution, 40 (3), msad035. 10.1093/molbev/msad035

