## Supplementary Information for "Museomics analyses inform about *Channichthys* icefish species diversity"

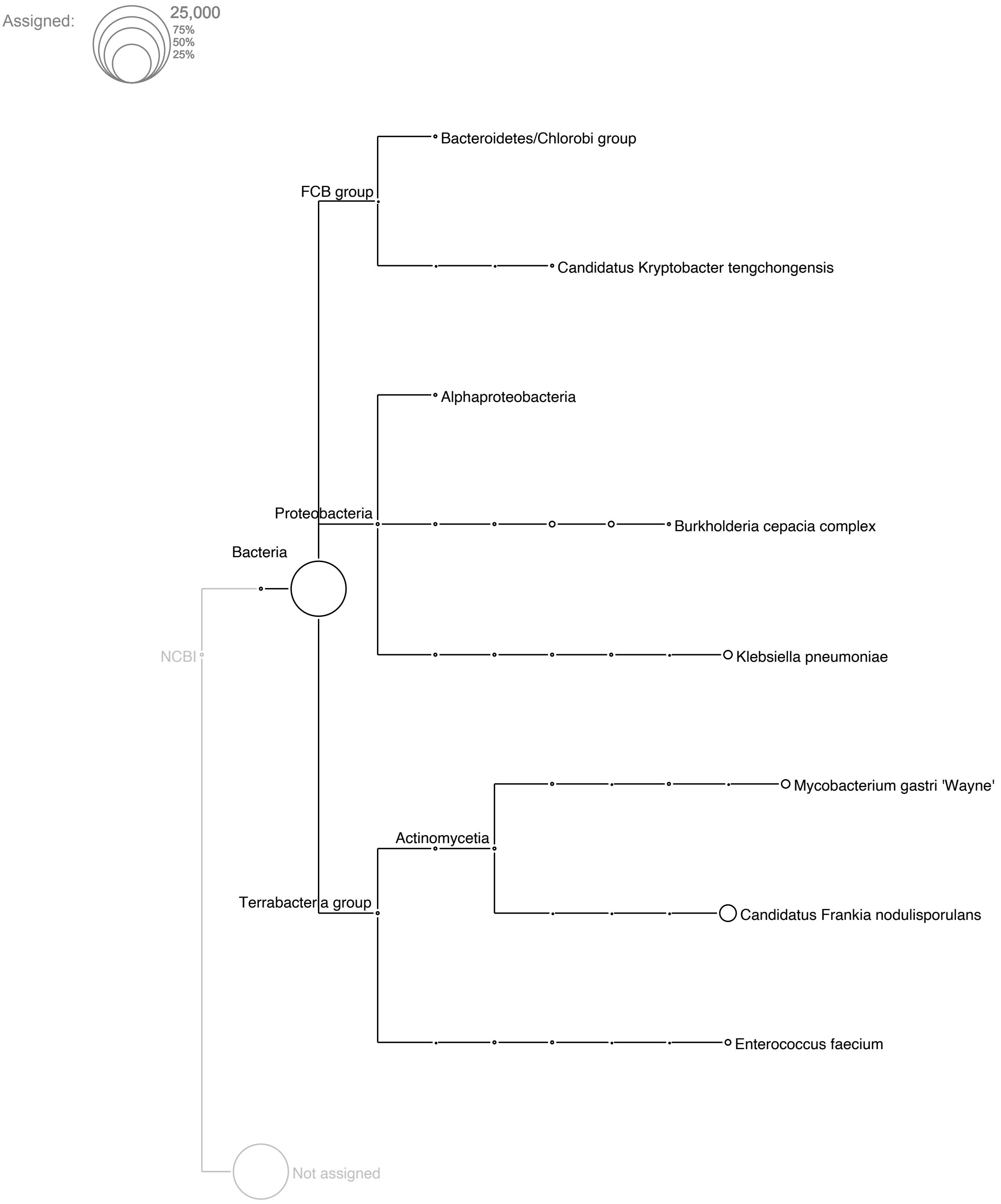


**Supplementary Figure 1** Taxonomic assignments of reads extracted from specimen ZIN 56635 (*C. rhinoceratus*). Circle size indicates numbers of reads assigned to a taxon.

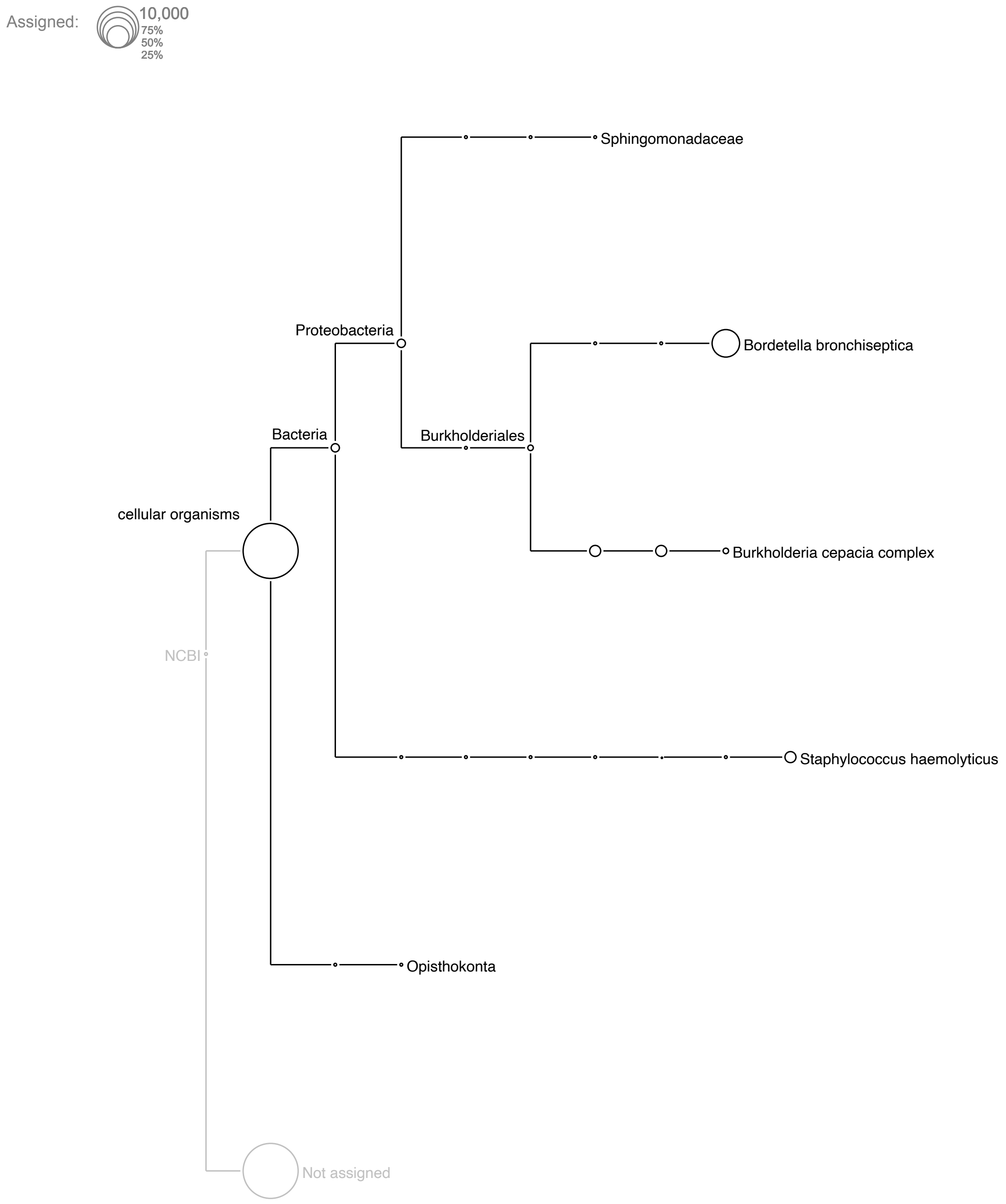


**Supplementary Figure 2** Taxonomic assignments of reads extracted from specimen ZIN 56636 (*C. rhinoceratus*). Circle size indicates numbers of reads assigned to a taxon.

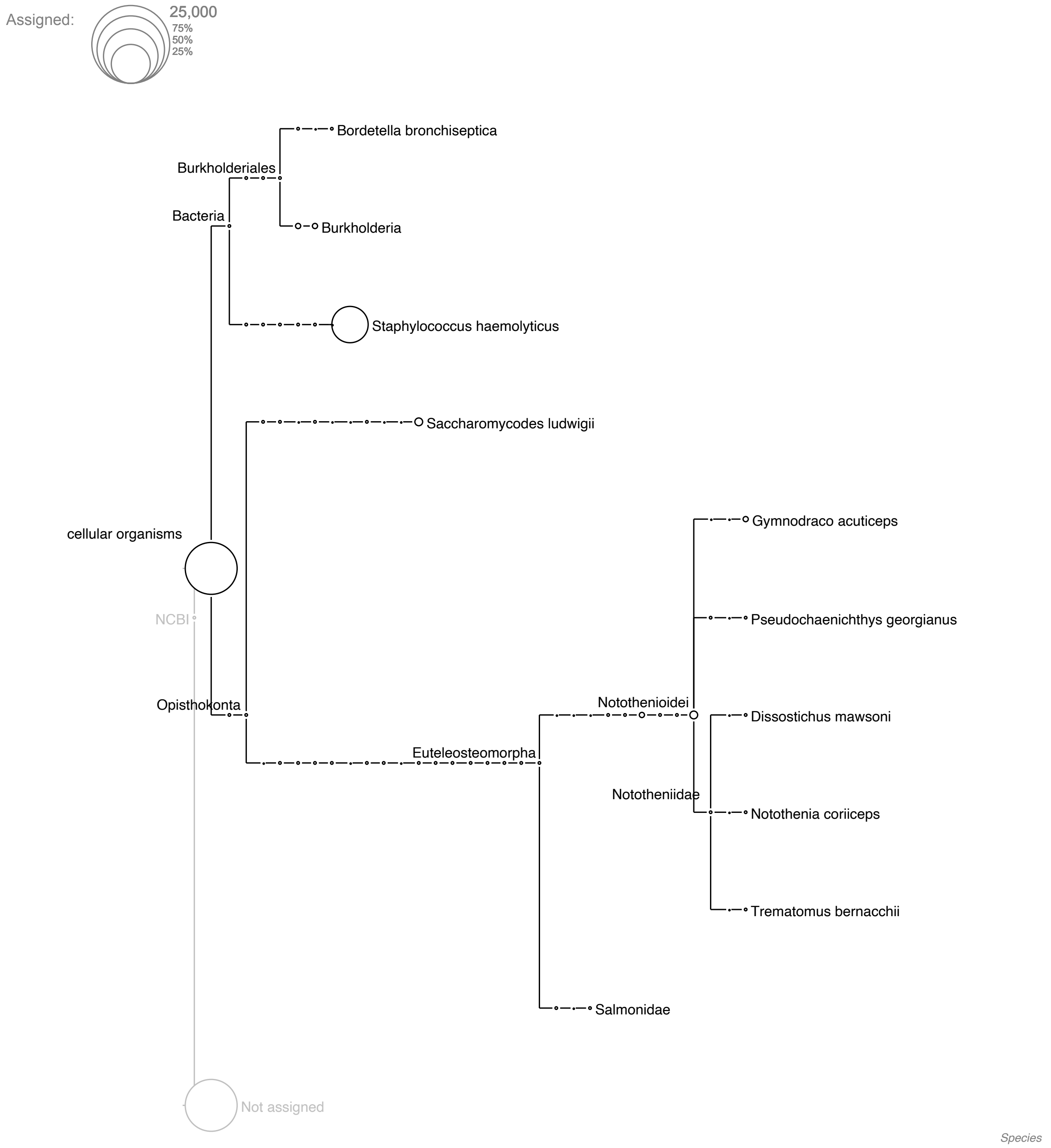


**Supplementary Figure 3** Taxonomic assignments of reads extracted from specimen ZIN 56638 (*C. rhinoceratus*). Circle size indicates numbers of reads assigned to a taxon.

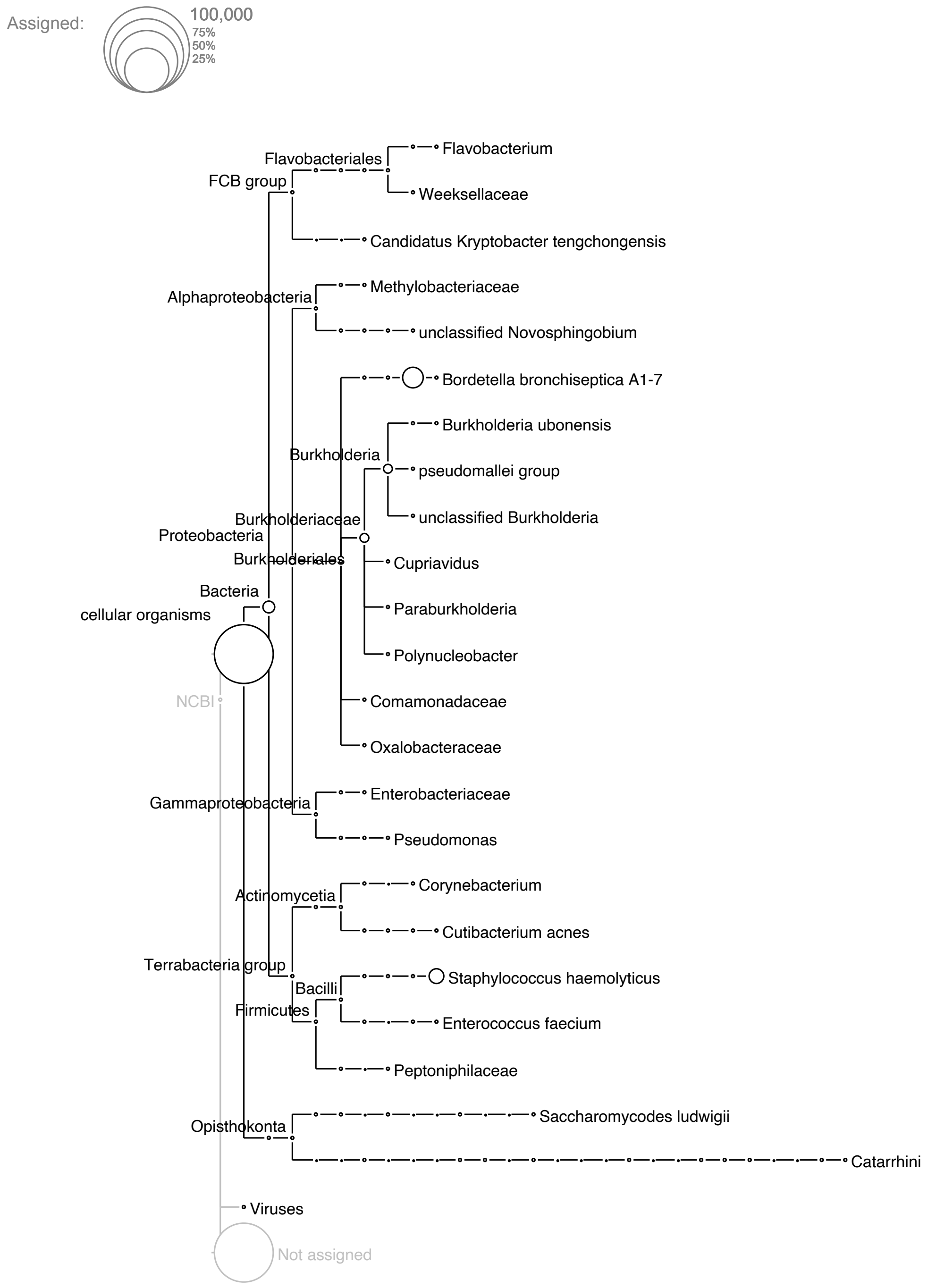


**Supplementary Figure 4** Taxonomic assignments of reads extracted from specimen ZIN 53007 (*C. rugosus*). Circle size indicates numbers of reads assigned to a taxon.

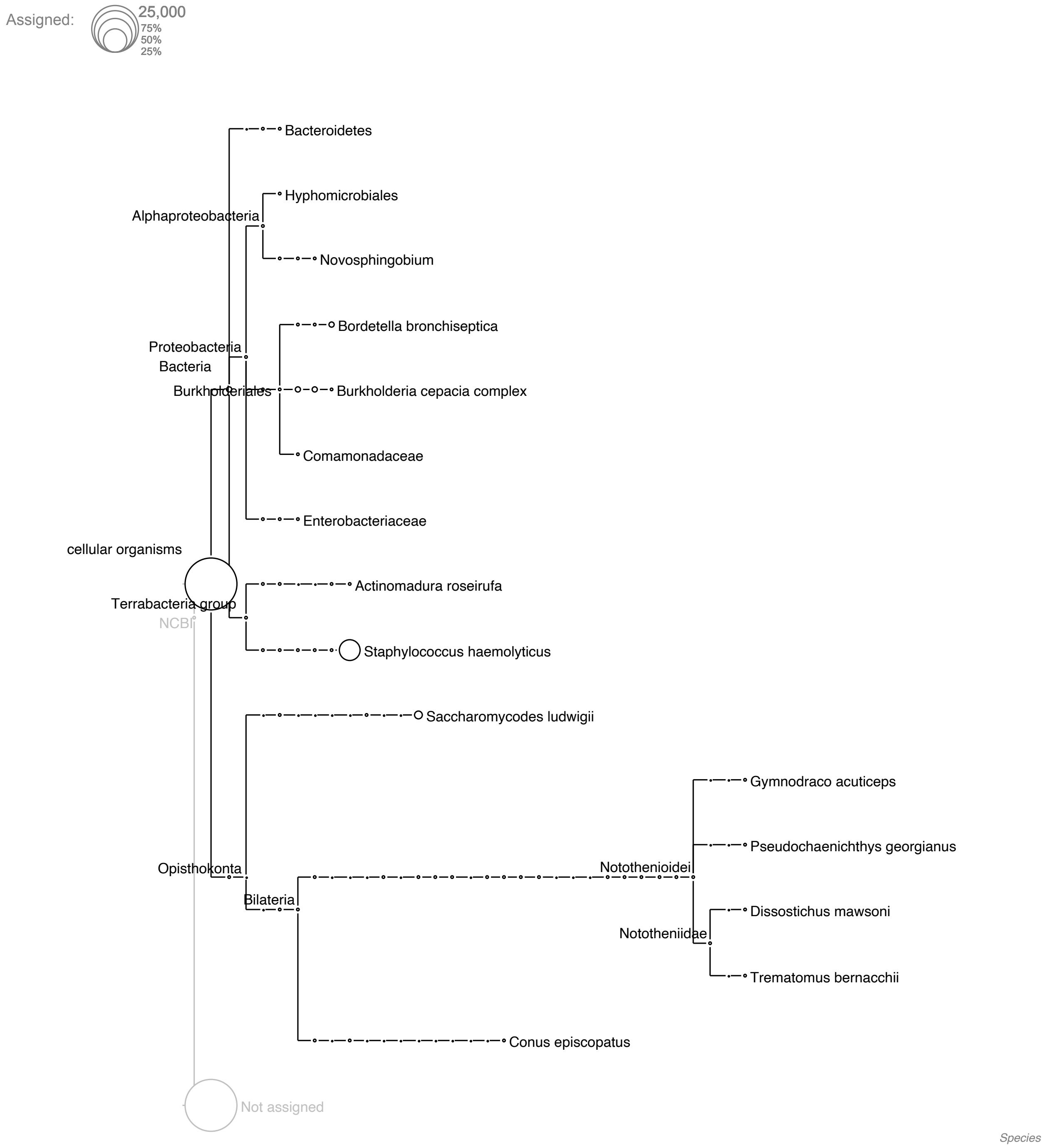


**Supplementary Figure 5** Taxonomic assignments of reads extracted from specimen ZIN 56294 (*C. rugosus*). Circle size indicates numbers of reads assigned to a taxon.

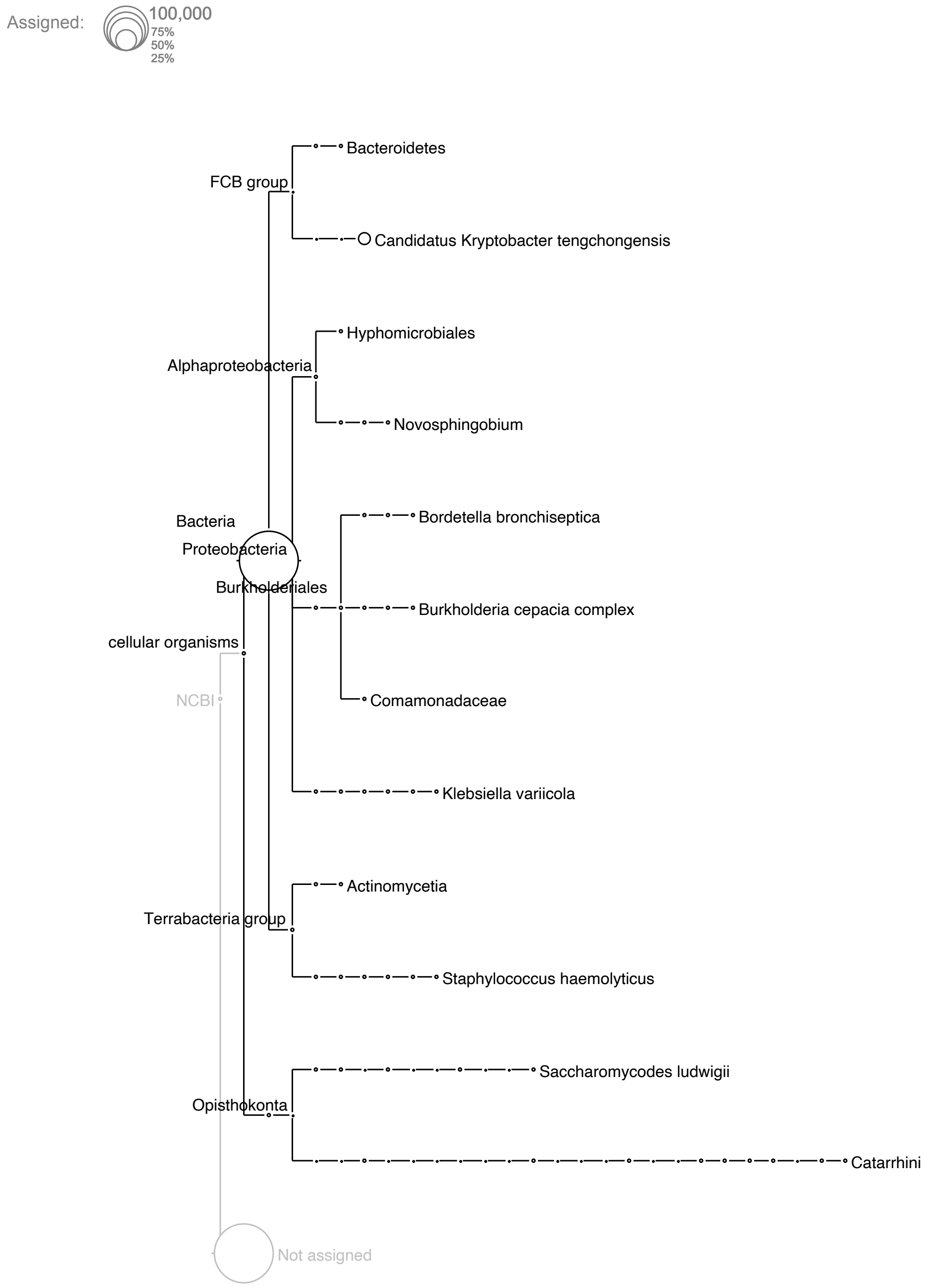


**Supplementary Figure 6** Taxonomic assignments of reads extracted from specimen ZIN 56292 (*C. rugosus*). Circle size indicates numbers of reads assigned to a taxon.

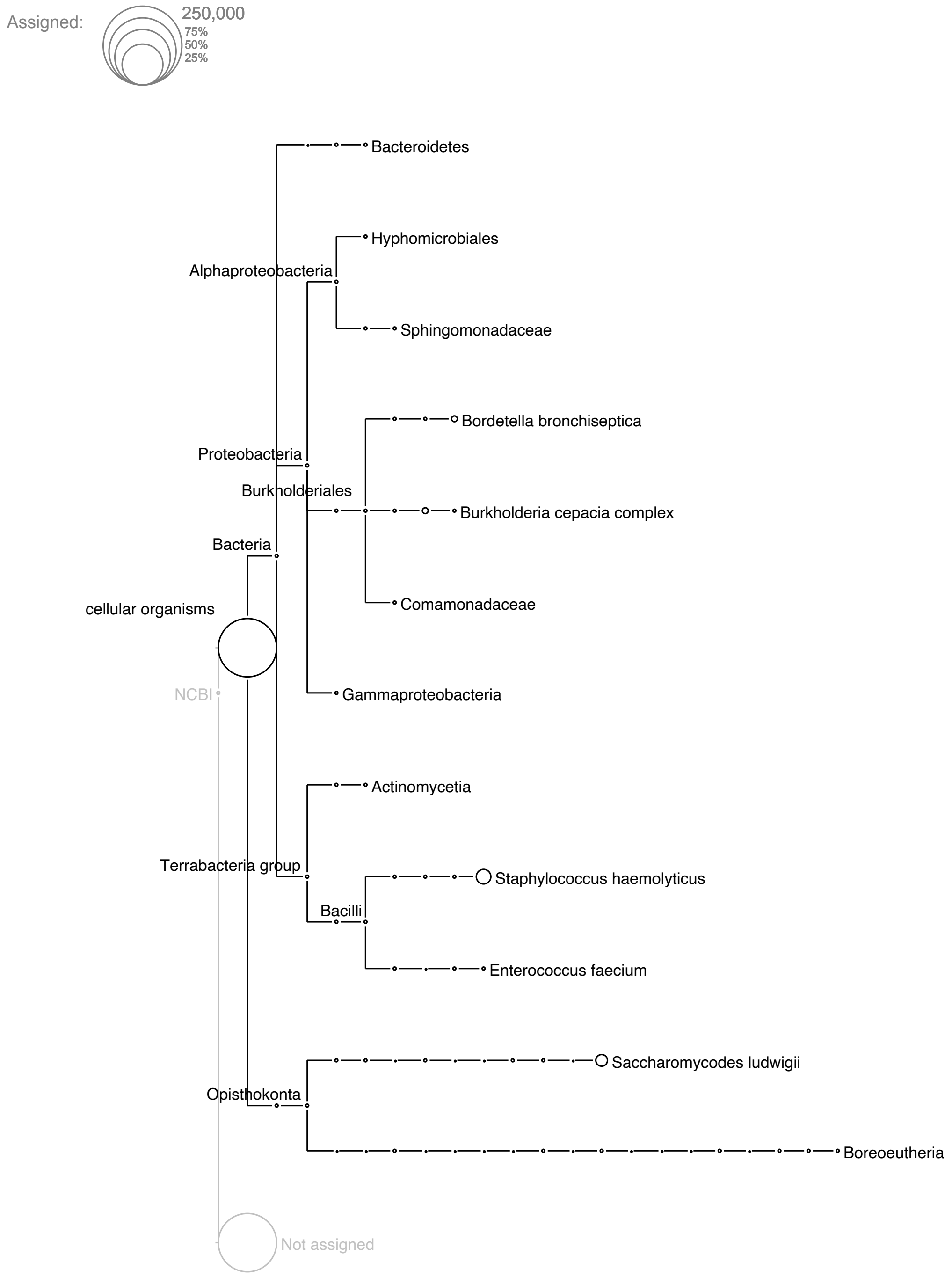


**Supplementary Figure 7** Taxonomic assignments of reads extracted from specimen ZIN 56273 (*C. velifer*). Circle size indicates numbers of reads assigned to a taxon.

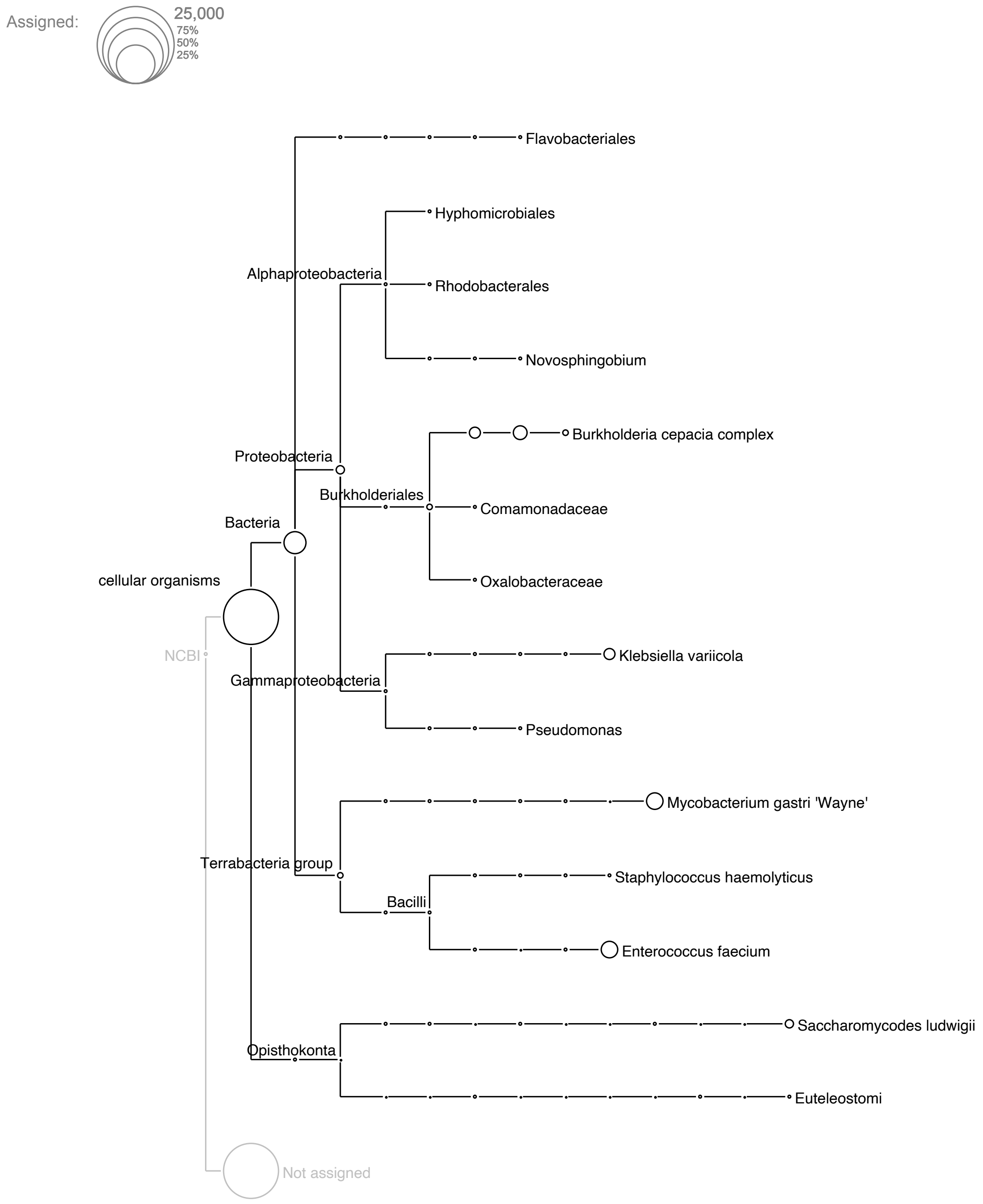


**Supplementary Figure 8** Taxonomic assignments of reads extracted from specimen ZIN 56274 (*C. velifer*). Circle size indicates numbers of reads assigned to a taxon.

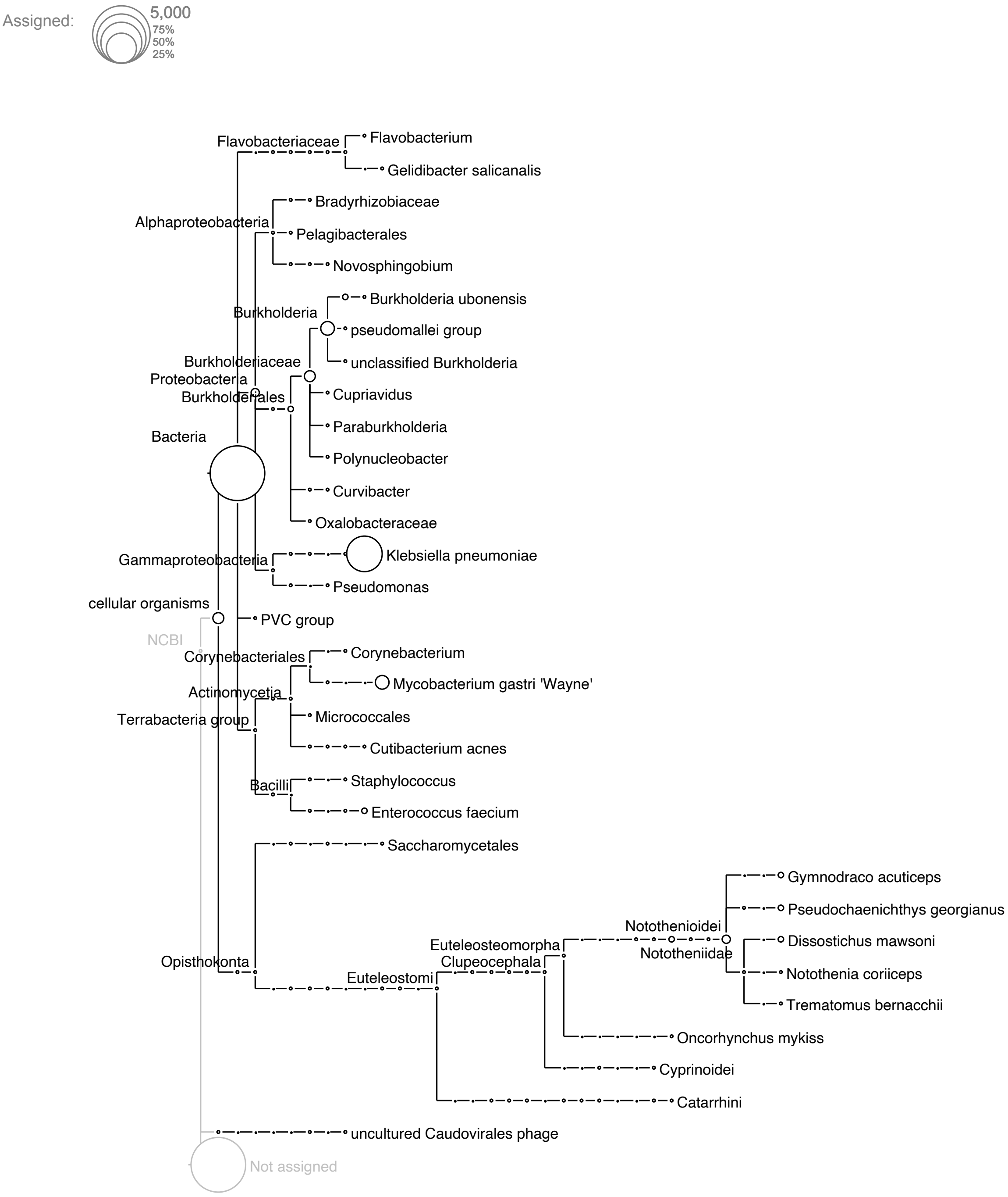


**Supplementary Figure 9** Taxonomic assignments of reads extracted from specimen ZIN 56275 (*C. velifer*). Circle size indicates numbers of reads assigned to a taxon.

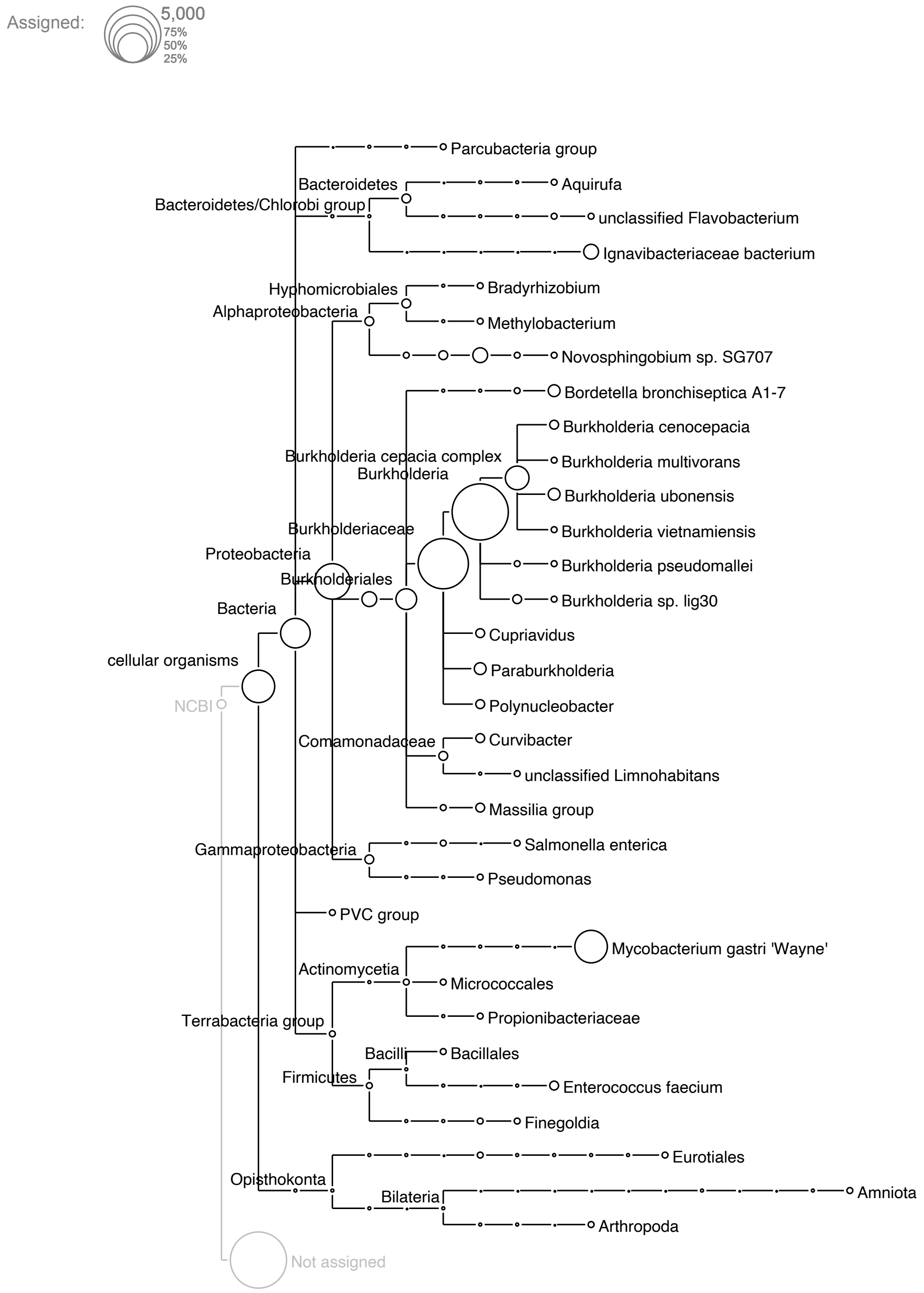


**Supplementary Figure 10** Taxonomic assignments of reads extracted from specimen ZIN 56536 (*C. panticapaei*). Circle size indicates numbers of reads assigned to a taxon.

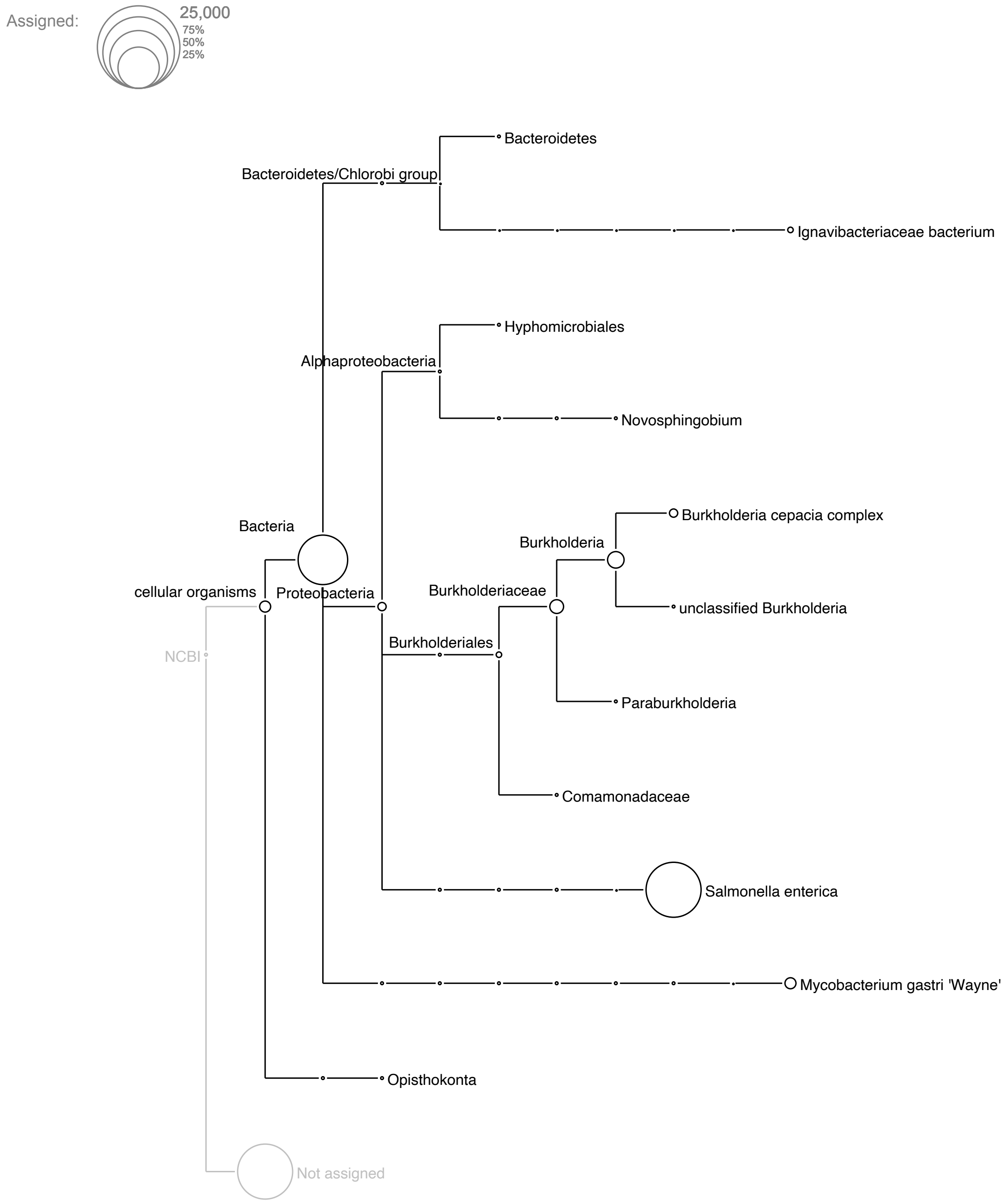


**Supplementary Figure 11** Taxonomic assignments of reads extracted from specimen ZIN 56537 (*C. panticapaei*). Circle size indicates numbers of reads assigned to a taxon.

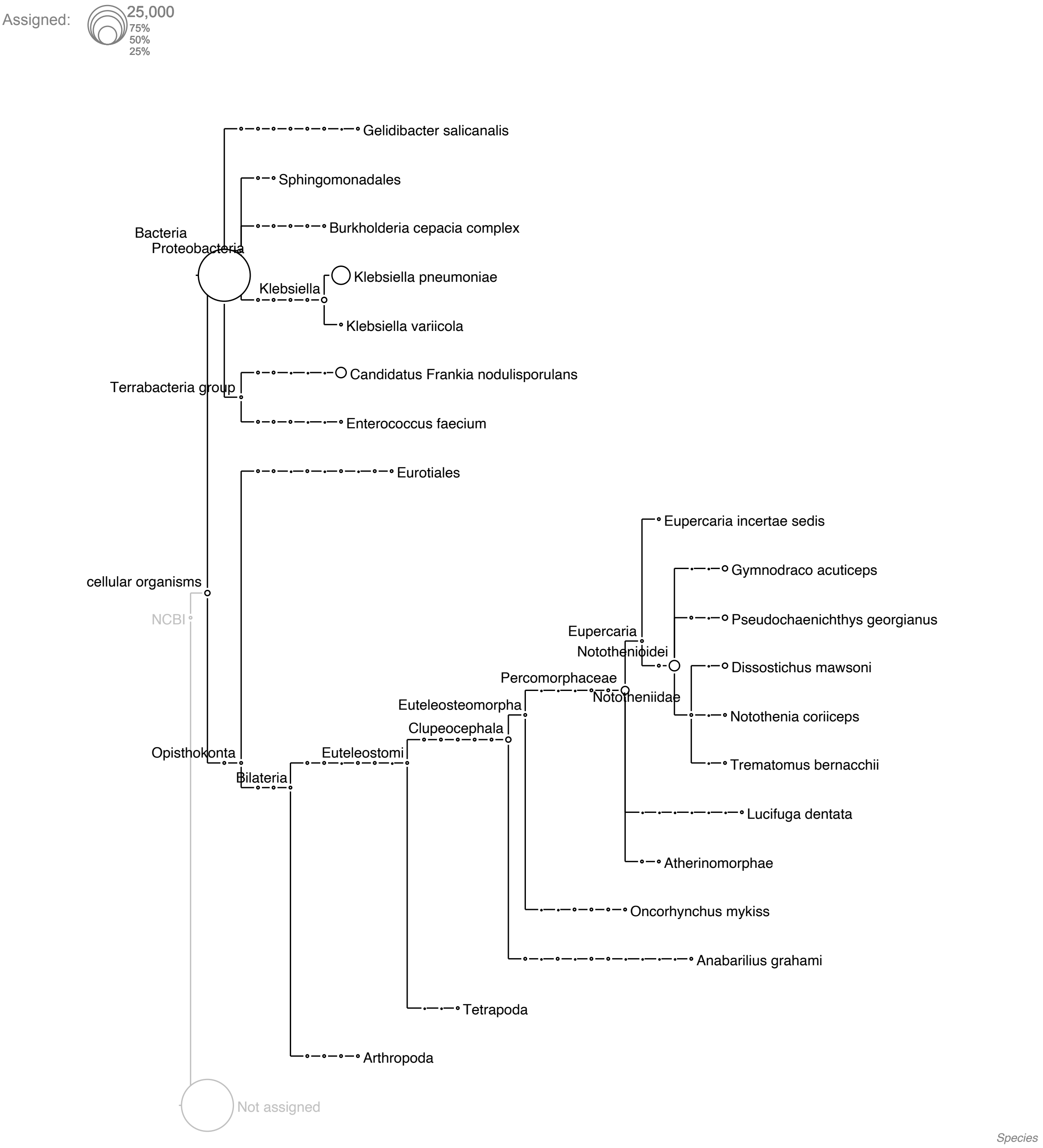


**Supplementary Figure 12** Taxonomic assignments of reads extracted from specimen ZIN 56538 (*C. panticapaei*). Circle size indicates numbers of reads assigned to a taxon.

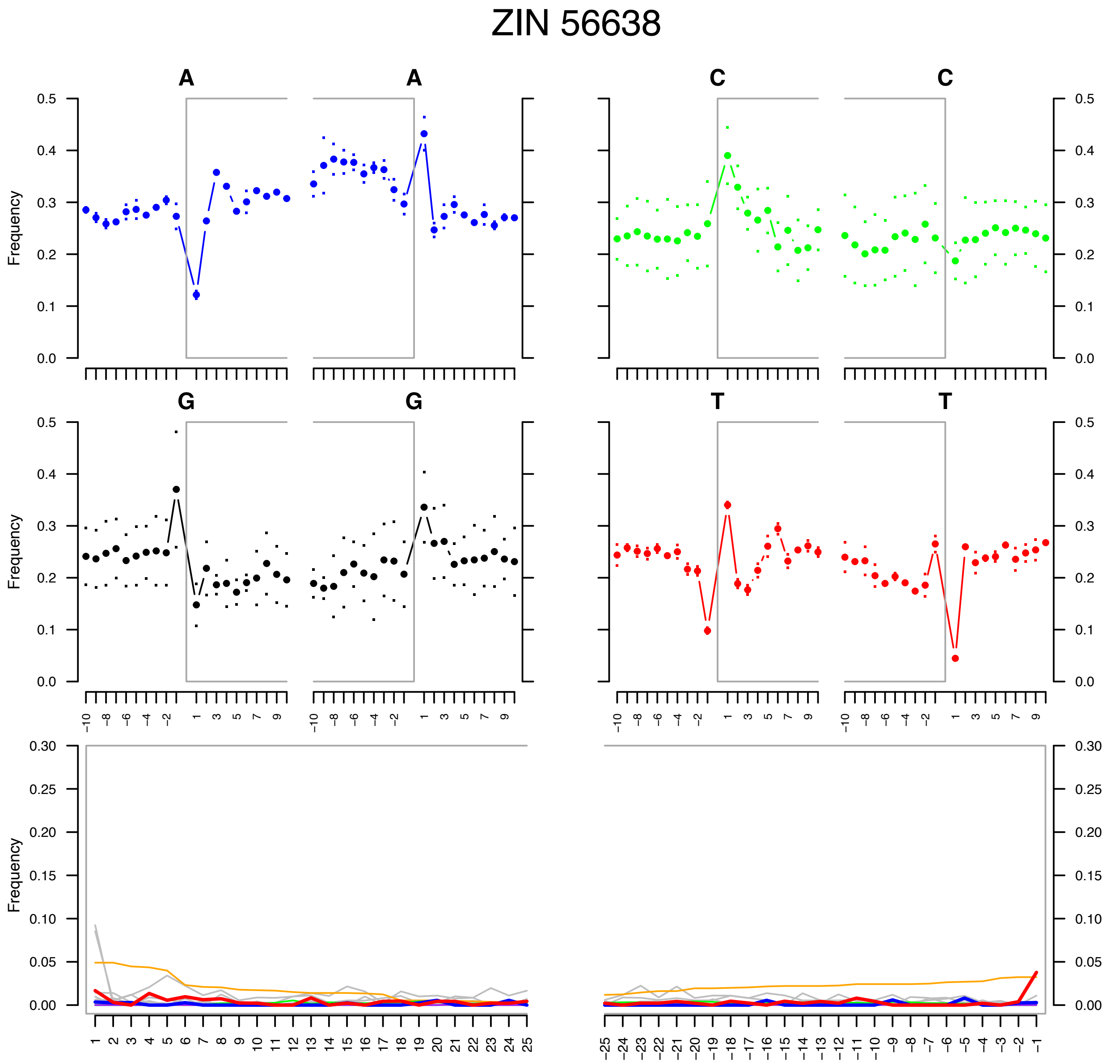


**Supplementary Figure 13** DNA damage patterns for reads extracted from specimen ZIN 56638 (*C. rhinoceratus*).

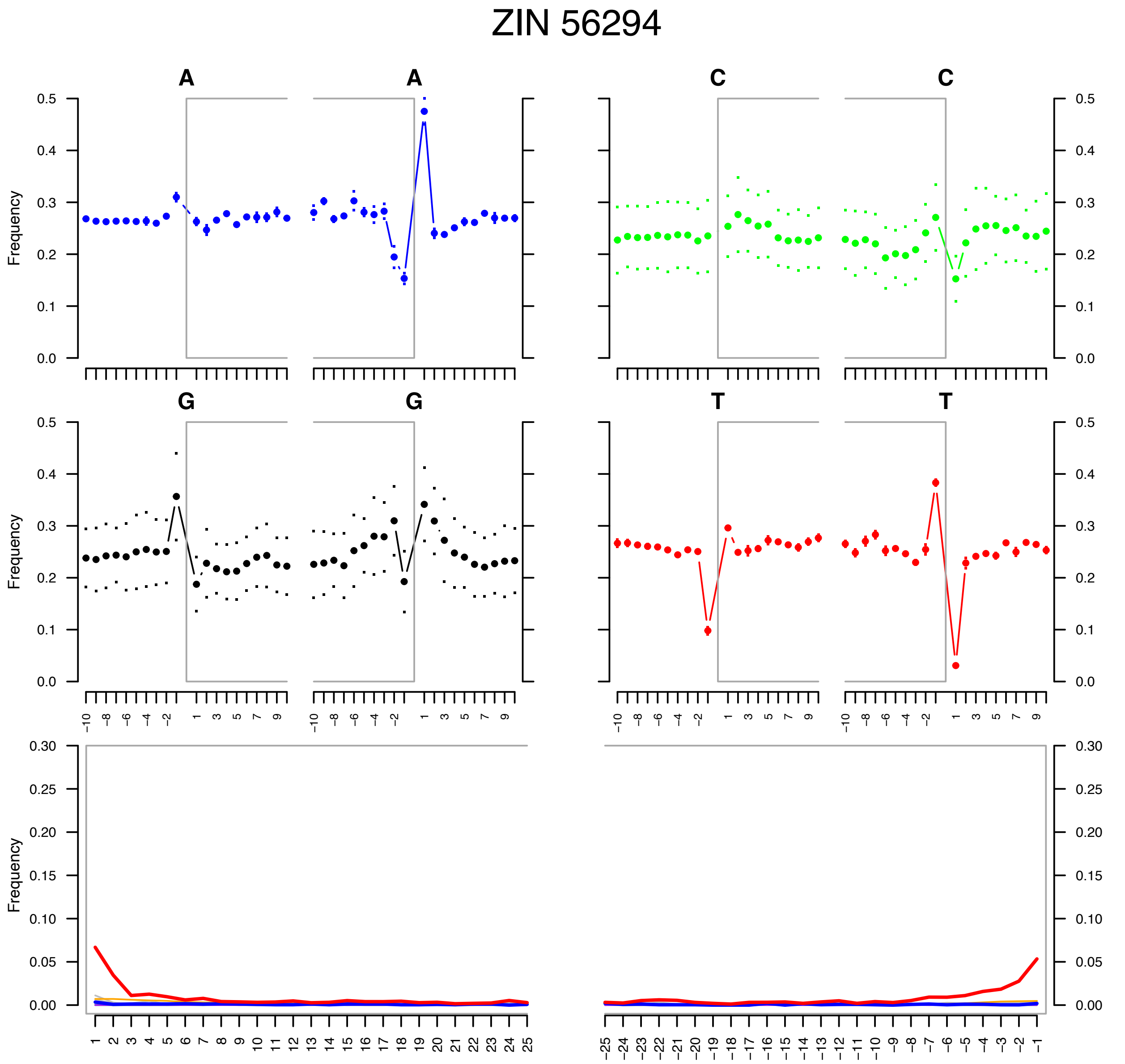


**Supplementary Figure 14** DNA damage patterns for reads extracted from specimen ZIN 56294 (*C. rugosus*).

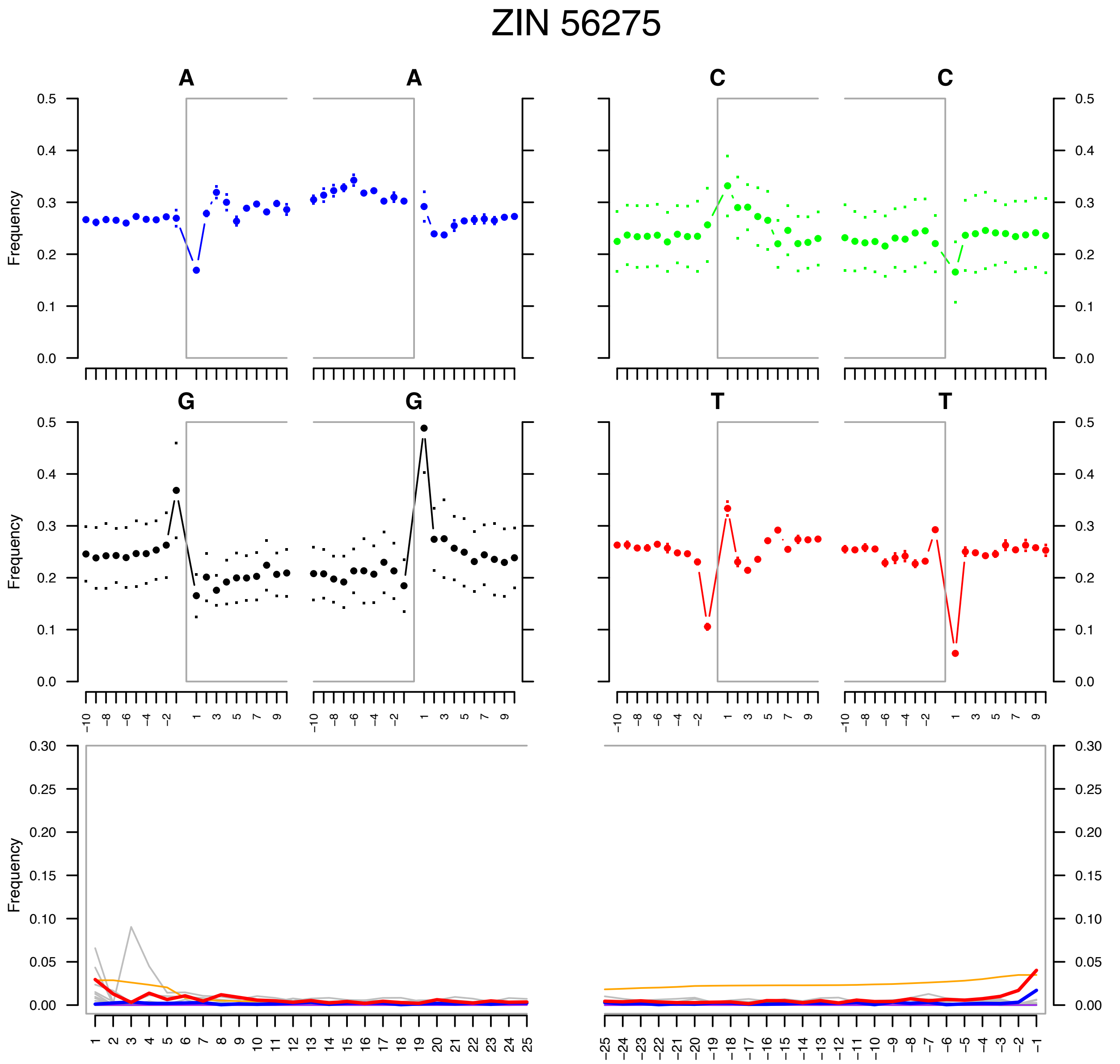


**Supplementary Figure 15** DNA damage patterns for reads extracted from specimen ZIN 56275 (*C. velifer*).

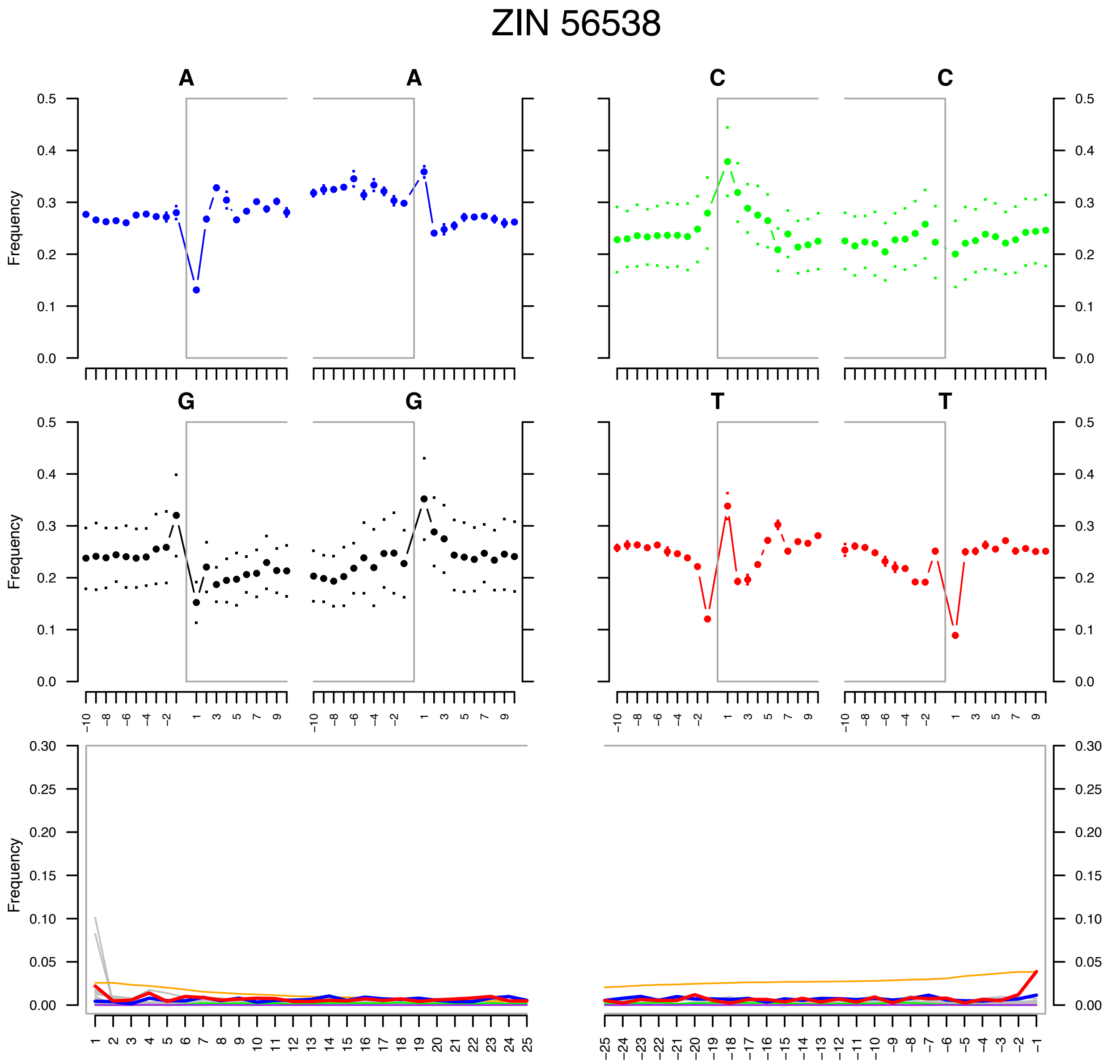


**Supplementary Figure 16** DNA damage patterns for reads extracted from specimen ZIN 56538 (*C. panticapaei*).

| Species | NCBI accession |
|---|---|
| ***Bovichtus angustifrons*** | MT559884.1 |
| ***Bovichtus argentinus*** | AP006020.1 |
| ***Eleginops maclovinus*** | KY038381.1 |
| ***Pleuragramma antarctica*** | NC_015652.1 |
| ***Aethotaxis mitopteryx*** | MT232658.1 |
| ***Dissostichus eleginoides*** | AB723627.1 |
| ***Dissostichus mawsoni*** | LC138011.1 |
| ***Lindbergichthys nudifrons*** | MT559890.1 |
| ***Pagothenia borchgrevinki*** | MT232659.1 |
| ***Trematomus tokarevi*** | MT559896.1 |
| ***Trematomus bernacchii*** | MN841276.1 |
| ***Trematomus eulepidotus*** | MT559895.1 |
| ***Trematomus lepidorhinus*** | MN864240.2 |
| ***Trematomus loennbergii*** | MT447073.1 |
| ***Notothenia coriiceps*** | JF933906.1 |
| ***Artedidraco skottsbergi*** | MT559883.1 |
| ***Dolloidraco longedorsalis*** | MT559886.1 |
| ***Pogonophryne albipinna*** | MN614417.1 |
| ***Akarotaxis nudiceps*** | MT559882.1 |
| ***Gymnodraco acuticeps*** | MT559888.1 |
| ***Parachaenichthys charcoti*** | KP300644.1 |
| ***Champsocephalus esox*** | NC_063099.1 |
| ***Champsocephalus gunnari*** | NC_018340.1 |
| ***Pagetopsis macropterus*** | MT559892.1 |
| ***Neopagetopsis ionah*** | MT559891.1 |
| ***Pseudochaenichthys georgianus*** | MT559893.1 |
| ***Chaenodraco wilsoni*** | MF536715.1 |
| ***Chionodraco myersi*** | DQ526430.1 |
| ***Chionodraco hamatus*** | KT921282.1 |
| ***Chionodraco rastrospinosus*** | MF622064.1 |
| ***Chaenocephalus aceratus*** | NC_015654.1 |
| ***Chionobathyscus dewitti*** | MN104591.1 |
| ***Cryodraco antarcticus*** | MK941847.1 |

**Supplementary Table 1** Notothenioid species and NCBI accession numbers included in the phylogenetic analysis based on coding gene sequences.
